# Defining the Characteristics of Type I Interferon Stimulated Genes: Insight from Expression Data and Machine Learning

**DOI:** 10.1101/2021.10.08.463622

**Authors:** Haiting Chai, Quan Gu, Joseph Hughes, David L. Robertson

## Abstract

A virus-infected cell triggers a signalling cascade resulting in the secretion of interferons (IFNs), which in turn induce the up-regulation of IFN-stimulated genes (ISGs) that play an important role in the inhibition of the viral infection and the return to cellular homeostasis. Here, we conduct detailed analyses on 7443 features relating to evolutionary conservation, nucleotide composition, gene expression, amino acid composition, and network properties to elucidate factors associated with the stimulation of genes in response to type I IFNs. Our results show that ISGs are less evolutionary conserved than genes that are not significantly stimulated in IFN experiments (non-ISGs). ISGs show significant depletion of GC-content in the coding region of their canonical transcripts, which leads to under-representation in the nucleotide compositions. Differences between ISGs and non-ISGs are also reflected in the properties of their coded amino acid sequence compositions. Network analyses show that ISG products tend to be involved in key paths but are away from hubs or bottlenecks of the human protein-protein interaction (PPI) network. Our analyses also show that interferon-repressed human genes (IRGs), which are down-regulated in the presence of IFNs, can have similar properties to ISGs, thus leading to false positives in ISG predictions. Based on these analyses, we design a machine learning framework integrating the usage of support vector machine (SVM) and feature selection algorithms. The ISG prediction achieves an area under the receiver operating characteristic curve (AUC) of 0.7455 and demonstrates the similarity between ISGs triggered by type I and III IFNs. Our machine learning model predicts a number of genes as potential ISGs that so far have shown no significant differential expression when stimulated with IFN in the cell types and tissue types compiled in the available IFN-related databases. A webserver implementing our method is accessible at http://isgpre.cvr.gla.ac.uk/.

**Author summary:** Interferons (IFNs) are signalling proteins secreted from host cells. IFN-triggered signalling activates the host immune system in response to intra-cellular infection. It results in the stimulation of many genes that have anti-pathogen roles in host defenses. Interferon-stimulated genes (ISGs) have unique properties that make them different from those not significantly up-regulated in response to IFNs (non-ISGs). We find the down-regulated interferon-repressed genes (IRGs) have some shared properties with ISGs. This increases the difficulty of distinguishing ISGs from non-ISGs. The use of machine learning is a sensible strategy to provide high throughput classifications of putative ISGs, for investigation with *in vivo* or *in vitro* experiments. Machine learning can also be applied to human genes for which there are insufficient expression levels before and after IFN treatment in various experiments. Additionally, the interferon type has some impact on ISG predictability. We expect that our study will provide new insight into better understanding the inherent characteristics of human genes that are related to response in the presence of IFNs.

## Introduction

Interferons (IFNs) are a family of cytokines originally defined for their capacity to interfere with viral replication. They are secreted from host cells after an infection by pathogens such as bacteria or viruses to trigger the innate immune response with the aim of inhibiting viral spread by ‘warning’ uninfected cells [1]. The response induced by IFNs is usually fast and feedforward, especially to synthesize new IFNs, which guarantees a full response even if the initial activation is limited [2]. In humans, seven IFNs have been discovered and grouped into three types based on distinct receptors on the cell surface, i.e., IFN-α receptor (IFNAR), IFN-γ receptor (IFNGR), and IFN-λ receptor 1 (IFNLR1)/interleukin-10 receptor 2 (IL-10R2) in the signalling cascade (Fig 1) [3]. Type I and III IFNs both help to regulate and activate host immune response, but the function of the latter group is less intense than the former [4–6]. Type II IFNs are also anti-pathogen, immunomodulatory, and proinflammatory but focus on establishing cell immunity [5, 7, 8]. All three types of IFNs are capable of activating the Janus kinase/signal transducer and activator of transcription (JAK-STAT) pathway and inducing the transcriptional up-regulation of approximately 10% of human genes that prime cells for stronger pathogen detections and defenses [9–11]. Henceforth, these up-regulated human genes are referred to as IFN-stimulated genes (ISGs). They play an important role in the establishment of cellular antiviral state, the inhibition of viral infection and the return to cellular homeostasis [8–10, 12]. For example, the ectopic expression of ISG: heparinase (HPSE) can inhibits the attachment of multiple viruses [13, 14]; interferon induced transmembrane proteins (IFITM) can impair the entry of multiple viruses and traffic viral particles to degradative lysosomes [15, 16]; MX dynamin like GTPase proteins (MX) can effectively block early steps of multiple viral replication cycles [17]. The qualitative nature of ISGs is determined by the balance between the activation of signal transducers and activators of transcription (STAT) and IFN-stimulated gene factor 3 complex (ISGF3)/IFN-γ activation factor (GAF) (Fig 1) [18]. Abnormality in the IFN-signalling cascade, e.g., the absence of signal transducer and activator of transcription 1 (STAT1) will lead to the failure of activating ISGs, making the host cell highly susceptible to virus infections [19].

**Fig 1.**
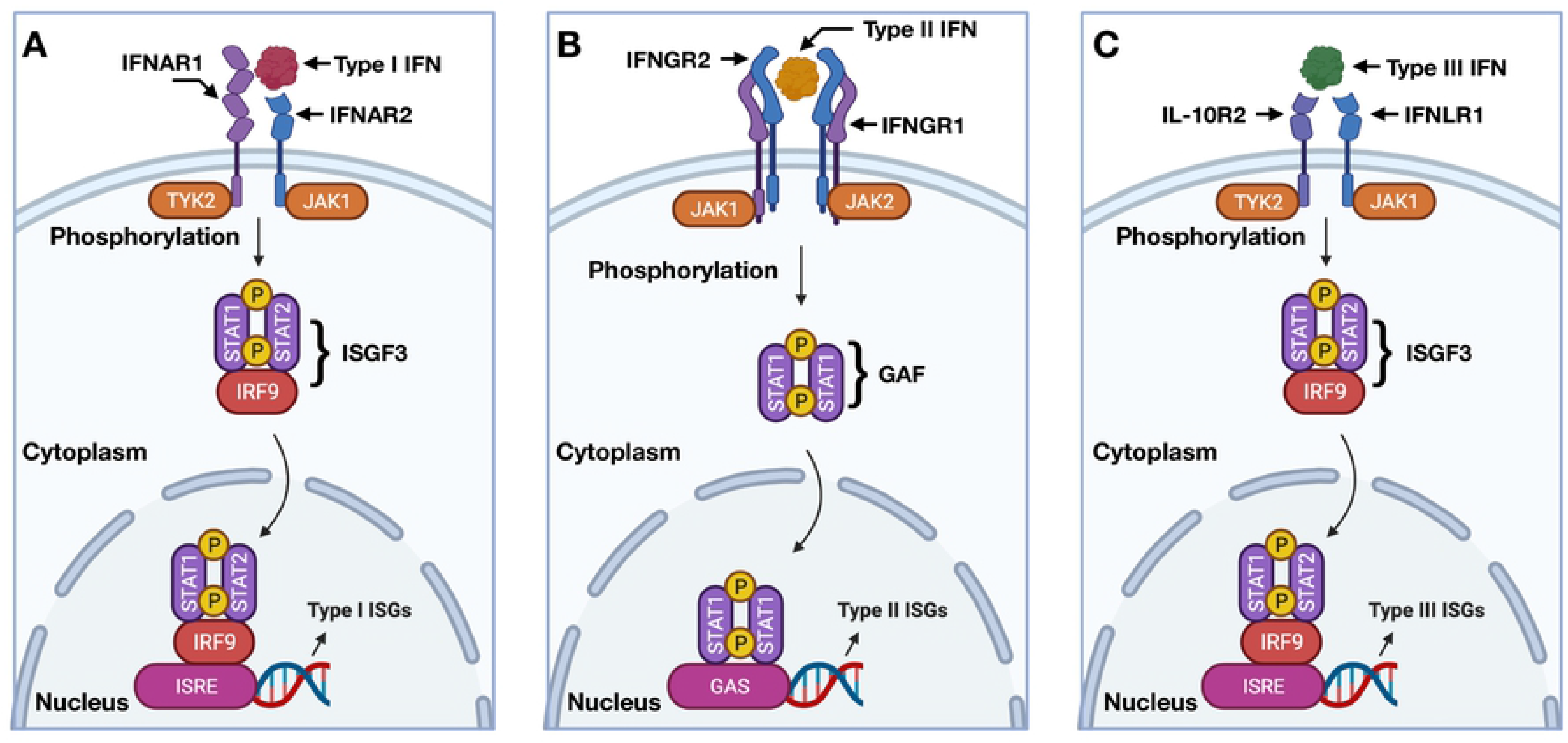
Illustration of signalling cascade triggered by different IFNs. In (A), type I IFN signals through IFNAR, Janus kinase 1 (JAK1), tyrosine kinase 2 (TYK2), STAT, and IFN-regulatory factor 9 (IRF9) to form ISGF3, and then bind to IFN-stimulated response elements (ISRE) to induce the expression of type I ISGs. In (B), type II IFN signals through IFNGR, JAK1 and JAK2 to form GAF and then bind to gamma-activated sequence promoter elements (GAS) to induce the expression of type II ISGs. In (C), type III IFN signals through IFNLR1, IL-10R2, JAK1, TYK2, STAT, and IRF9 to form ISGF3, and then bind to ISRE to induce the expression of type III ISGs. Figure created using the BioRender (https://biorender.com/). Abbreviations: IFNs, interferons; IFNAR, IFN-α receptor; ISGF3, IFN-stimulated gene factor 3 complex; ISGs, interferon-stimulated human genes; IFNGR, IFN-γ receptor; GAF, IFN-γ activation factor; IFNLR1, IFN-λ receptor 1; IL-10R2, interleukin-10 receptor 2; STAT, signal transducers and activators of transcription.

Most research on ISGs has focused on elucidating the role of ISGs in antiviral activities or discovering new ISGs within or across species [8-10, 15, 20, 21]. The identification of ISGs can be achieved via various approaches. Associating gene expression with suppression of viral infection is a good strategy to identify ISGs with obvious antiviral performance, exemplified by the influenza inhibitor, MX dynamin like GTPase 1 (MX1), and the human immunodeficiency virus 1 inhibitor, MX dynamin like GTPase 2 (MX2) [17]. CRISPR screening is a loss-of-function experimental approach to identify ISGs required for IFN-mediated inhibition to viruses, e.g., it enabled the discovery of tripartite motif containing 5 (TRIM5), MX2 and bone marrow stromal cell antigen 2 (BST2) [22]. Monitoring the ectopic expression of ISGs is an instrumental way to find some ISGs that are individually sufficient for viral suppression, e.g. interferon stimulated exonuclease gene 20 (ISG20) and ISG15 ubiquitin like modifier (ISG15) [23]. Using fold change-based criterion to measure whether a target human gene is induced by IFN signalling now has become a well-accepted idea, but the upregulation cut-off may vary in different studies [21, 24, 25]. The online database, Interferome (http://www.interferome.org), provides an excellent resource by compiling *in vivo* and *in vitro* gene expression profiles in the context of IFN stimulation [21]. The Orthologous Clusters of Interferon-stimulated Genes (OCISG, http://isg.data.cvr.ac.uk) provides an evolutionary comparative approach of genes differentially expressed in the type I IFN system for ten different species [8]. The later study employed a standardised experimental protocol of fibroblast cells stimulated by type I IFN.

Although these studies contribute to a better understanding and detection of ISGs, the knowledge they compiled was limited to a specific IFN type in specific organs, tissues or cells [2]. Despite some well-investigated ISGs, the majority of classified ISGs have limitedly expression following IFN stimulations [8, 21], which means the difference between ISGs and those human genes not significantly up-regulated in the presence of IFNs (non-ISGs) may not be obvious especially when being assessed more generally. It should also be noted that, within non-ISGs, there are a group of genes down-regulated during IFN stimulations. Here, we refer to them as interferon-repressed human genes (IRGs) and they constitute another major part of the IFN regulation system [8, 26]. Collectively, the complex nature of the IFN-stimulated system results in knowledge that is far from comprehensive.

Hence, we seek to characterise the properties of ISGs and to determine whether genes can be identified as ISGs using an *in-silico* machine learning approach. We choose experimental data from human fibroblast cells as the baseline and focus on human genes stimulated by the type I IFNs. We construct a refined high-confidence dataset consisting of 620 ISGs and 874 non-ISGs by cross-checking the genes across multiple databases including the OCISG [8], Interferome [21], and Reference Sequence (RefSeq) [27]. The analyses are conducted primarily on our refined data using genome-and proteome-based features that are likely to influence the expression of human genes in the presence of type I IFNs. Then based on the calculated features, we design a machine learning framework with an optimised feature selection strategy for the prediction of putative ISGs in different IFN systems. Finally, we also develop an online webserver that implements our machine learning method at http://isgpre.cvr.gla.ac.uk/.

## Methods

### Dataset preparation

In this study, we retrieve 2054 ISGs (Log_2_(Fold Change)>2), 12379 non-ISGs (Log_2_(Fold Change) <1), and 3944 human genes with low expression levels in IFN experiments (ELGs, expression-limited genes with less than 1 count per million reads mapping across the three biological replicates [28, 29]) from the OCISG (http://isg.data.cvr.ac.uk/) [8]. Gene clusters in the OCISG are built by using the Ensembl Compara database [30], which provides a thorough account of gene orthology based on whole genomes available in the Ensembl database [31]. Labels of these human genes are defined based on the fold change (before and after IFN treatments) and a false discovery rate following IFN treatments in human fibroblast cells. We search the collected 18377 entries against the RefSeq database (https://www.ncbi.nlm.nih.gov/refseq/) [27] to decipher features based on appropriate transcripts (canonical) [32] coding for the main functional isoforms of these human genes, obtaining 1315, 7304, and 2217 results for ISGs, non-ISGs and ELGs, respectively. These 10836 human genes are well-annotated by multiple online databases and are used as the background set (i.e., dataset S1) in the analyses.

For the purpose of generating a set of human genes with high confidence of being interferon-up-regulated and non-up-regulated in response to the type I IFNs, we search labelled human genes against the Interferome database (http://www.interferome.org/) [21]. We filter out ISGs without high up-regulation (Log_2_(Fold Change) > 1.0) or with obvious down-regulation (Log_2_(Fold Change) < −1.0) in the presence of type I IFNs. This procedure guarantees a refined ISGs dataset with strong levels of stimulation induced by type I IFNs and reduces biases driven by IRGs for the analyses and predictions. We filter out non-ISGs showing enhanced expression after type I IFN treatments (Log_2_(Fold Change) > 0). The exclusion of these non-ISGs can effectively reduce the risk of involving false negatives in analyses and producing false positives in predictions. As a result, the refined dataset S2 contains 620 ISGs and 874 non-ISGs with relatively high confidence.

The training procedure in the machine learning framework is conducted on a balanced dataset: S2’ consisted of 992 randomly selected ISGs and non-ISGs from dataset S2. The remaining human genes in S2 are used for independent testing. Additionally, we also construct another six testing datasets for the purpose of review and assessment. Dataset S3 contains 695 ISGs with low confidence compared to those ISGs in dataset S2. Some of them could be IRGs in the type I IFN system. Dataset S4 contains 1006 IRGs from the human fibroblast cell experiments. Dataset S5, S6, and S7 are constructed based on records for experiments in type I, II, and III IFN systems from the Interferome database [21]. The criterion for an ISG in the latter three datasets is a high level of up-regulation (Log_2_(Fold Change) > 1.0) while that for non-ISGs is no up-regulation after IFN treatments (Log_2_(Fold Change) < 0). The last testing dataset S8 is derived from our background dataset S1, containing 2217 ELGs. A breakdown of the aforementioned eight datasets is shown in Table 1. Detailed information of the human genes used in this study is provided in **S1 Data**.

**Table 1.** A breakdown of datasets used in this study.

| Dataset | Brief description | IFN system | ISGs | Non-ISGs | ELGs |
| --- | --- | --- | --- | --- | --- |
| S1 | Well-annotated human genes | Type I in fibroblast cells | 1315 | 7304 | 2217 |
| S2 | Refined dataset with high confidence | Type I in fibroblast cells | 620 | 874 | 0 |
| S2' | Training subset of S2 | Type I in fibroblast cells | 496 | 496 | 0 |
| S2'' | Testing subset of S2 | Type I in fibroblast cells | 124 | 378 | 0 |
| S3 | ISGs with low confidence in S1 | Type I in fibroblast cells | 695 | 0 | 0 |
| S4 | IRGs divided from S1 | Type I in fibroblast cells | 0 | 1006 | 0 |
| S5 | ISGs from the Interferome database [21] | Type I in all cells | 1259 | 872 | 0 |
| S6 | ISGs from the Interferome database [21] | Type II in all cells | 2229 | 755 | 0 |
| S7 | ISGs from the Interferome database [21] | Type III in all cells | 33 | 1683 | 0 |
| S8 | ELGs divided from S1 | Type I in fibroblast cells | 0 | 0 | 2217 |

**Abbreviations:** ISGs, interferon-stimulated human genes; non-ISGs, human genes not significantly up-regulated by interferons; IRGs, interferon-repressed human genes, ELGs, human genes with limited expression in interferon experiments.

### Generation of parametric features

We encode 397 parametric features from aspects of evolution, nucleotide composition, transcription, amino acid composition, and network preference. From the perspective of evolution, we use the number of transcripts and open reading frames (ORFs) to reflect alternative splicing diversity and gene polymorphism respectively. Genes with more transcripts and ORFs have higher alternative splicing diversity and polymorphism to produce proteins with similar or different biological functions [33, 34]. We use the number of protein-coding exons in the canonical transcripts to reflect the complexity of the alternative splicing [35]. Genes with more protein-coding exons in their canonical transcripts are considered to produce more complex alternative splicing products [36]. Here, duplication and mutation features are measured by the number of within species paralogues and substitutions [37, 38]. These data are collected from the BioMart [31] to assess the selection on protein sequences and mutational processes affecting the human genome [39].

From the perspective of nucleotide composition, we calculate the percent of adenine, thymine, cytosine, guanine, and their four-category combinations in the coding region of the canonical transcript. The first category measures the proportion of two different nitrogenous bases out of the implied four bases, e.g., GC-content. The second category also focuses on the combination of two nucleotides but involves the impact of phosphodiester bonds along the 5’ to 3’ direction, e.g., CpG-content [40]. The third category calculates the occurrence frequency of 4-mers, e.g., ‘CGCG’ composition to involve some positional resolution [41]. The last category considers the co-occurrence of some short linear motifs (SLims) in the complementary DNA (SLim_DNAs). From the perspective of transcription, we calculate the usage of 61 coding codons and three stop codons in the coding region of the canonical transcripts. Codon usage biases are observed when there are multiple codons available for coding one specific amino acid. They can affect the dynamics of translation thus regulate the efficiency of translation and even the folding of the proteins [42, 43].

From the perspective of amino acid composition, we calculate the percentage of 20 standard amino acids and their combinations based on their physicochemical properties [44]. Patterns in the amino acid level are considered to have a direct impact on the establishment of biological functions or to reflect the result of strong purifying selection [45]. Based on the chemical properties of the side chain, we group amino acids into seven classes including aliphatic, aromatic, sulfur, hydroxyl, acidic, amide, and basic amino acids. We also group amino acids based on geometric volume, hydropathy, charge status, and polarity, but find some overlaps among these features. For instance, amino acids with basic side chains are all positively charged. Aromatic amino acids all have large geometric volumes (volume > 180 cubic angstroms). Likewise, we also consider the co-occurrence of some SLims at the protein level. These co-occurring SLims in the protein sequence (SLim_AAs) may relate to potential mechanisms regulating the expression of ISGs [46].

When trying to measure the network preference for the gene products, we construct a human protein-protein interaction (PPI) network based on 332,698 experimentally verified interactions (confidence score > 0.63) from the Human Integrated Protein-Protein Interaction rEference database (HIPPIE, http://cbdm-01.zdv.uni-mainz.de/~mschaefer/hippie/) [47]. Nodes and edges of this network are provided at http://isgpre.cvr.gla.ac.uk/. Eight network-based features including the average shortest path, closeness, betweenness, stress, degree, neighbourhood connectivity, clustering coefficient, and topological coefficient are calculated from this network. Isolated nodes or proteins are not included in our network and are assigned zero value for all these eight features. The shortest path measures the average length of the shortest path between a focused node and others in the network. Closeness of a node is defined as the reciprocal of the length of the average shortest path. Proteins with a low value of the shortest paths or closeness are close to the centre of the network. Betweenness reflects the degree of control that one node exerted over the interactions of other nodes in the network [48]. Stress of a node measures the number of shortest paths passing through it. Proteins with a high value of betweenness or stress are close to the bottleneck of the network. Degree of a node counts the number of edges linked to it while neighbourhood connectivity reflected the average degree of its neighbours. Proteins with high degree or neighbourhood connectivity are close to the hub of the network. They are considered to play an important role in the establishment of the stable structure of the human interactome [49]. Clustering and topological coefficient measure the possibility of a node to form clusters or topological structures with shared neighbours. The former coefficient can be used to identify the modular organisation of metabolic networks [50] while the latter one may be helpful to find out virus mimicry targets [51].

### Generation of non-parametric features

In this study, non-parametric features are used to check the occurrence of SLims in the genome and proteome. The SLim_DNAs we constructed in this study contain three to five random nucleotides, producing 708,540 alternative choices. SLim_DNAs with no restrictions on their first or last position are not taken into consideration as their patterns can be expressed in a more concise way. A SLim_DNA will be picked out to encode a binary feature when its occurrence level in the coding region of the canonical ISG transcripts is significantly higher than that for non-ISGs (Pearson’s chi-squared test: *p* < 0.05). SLim_AAs are constructed with three to four fixed amino acids separated by putative gaps. The gap can be occupied by at most one random amino acid, producing 1,312,000 alternative choices. Likewise, binary features are prepared for SLim_AAs showing significant enrichment in ISGs products than in non-ISG products (Pearson’s chi-squared test: *p* < 0.05). Since there are lots of results rejecting the null-hypothesis, we adopt the Benjamini-Hochberg correction procedure to avoid type I error [52]. Additionally, we also encode two features to check the co-occurrence or absence of multiple SLim_DNAs and SLim_AAs. This co-occurrence status may be a better representation of functional sites composed of short stretches of adjacent nucleobases or amino acids surrounding SLim_DNAs or SLim_AAs[45]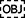.

### Assessment of associations between feature representation and IFN-triggered stimulations

In this study, we obtain 8619 human genes with expression data from the OCISG [8]. 4111 of them are annotated with a positive Log_2_(Fold Change) ranging from 0 to 12.6, which means they are up-regulated after IFN treatments. In order to measure the average level of feature representation (AREP) for genes with similar expression during IFN stimulations, we introduce a 0.1-length sliding-window to divide the data into 126 bins with different Log_2_(Fold Change). Here, Pearson’s correlation coefficient (PCC) is introduced to test the association between the representation of parametric features and IFN-triggered stimulation (Log_2_(Fold Change) > 0). It can be formulated as:

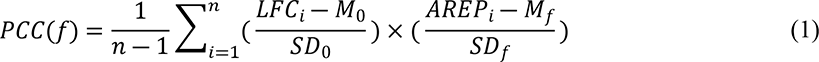

where *n* is the number of divided parts that equals to 126 in this study; *LFC*_*i*_ and *AREP*_*i*_ are the value of Log_2_(Fold Change) and AREP in the *i*-th part; *M*_0_ and *SD*_0_ are the mean and standard deviation of Log_2_(Fold Change), which is set as 6.4 and 3.7 respectively in this study; *M*_*f*_ and *SD*_*f*_ are the mean and standard deviation of 126 AREP that reflect the representation of the considered feature. To make fair comparisons among features with different scales, we normalise them based on the major value of their representations:

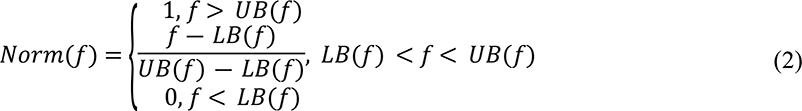

where *LB*(*f*) and *UB*(*f*) are the lower and upper bound representing the 5th and 95th percentile within representation values for the target feature. The representation of feature is considered to have a stronger positive/negative association with IFN-triggered stimulations if the PCC calculated from the normalised features is closer to 1.0/-1.0 and the p value calculated by the Student t-test is lower than 0.05.

### Machine learning and optimisation

In this study, we introduce a machine learning framework for the prediction of ISGs. Firstly, all features are encoded and normalised based on their major representations (**Equation 2**). Then we use an under-sampling procedure to generate a balanced dataset from the main dataset for training and modelling. Support vector machine (SVM) with radial basis function (RBF) [53] is used as the basic classifier, and it maps the normalised feature space to a higher dimension to generate a space plane to better classify the majority of positive and negative samples. Since there are usually lots of noisy data distributed in the feature space, it is necessary to remove disruptive features. This will effectively reduce the dimensionality of the feature space and make it easier for the SVM model to generate a more appropriate classification plane that involved fewer false positives and false negatives. Here, we develop a subtractive iteration algorithm driven by the change of area under the receiver operating characteristic curve (AUC) to filter out disruptive features (Fig 2). In each iteration, we traverse the features and remove those that do not improve the AUC of the prediction results. Theoretically, this algorithm can greatly optimise the feature space and remove all disruptive features after multiple iterations. In the testing procedure, we encode the optimum features for testing samples and place them in the optimised feature space. Samples with longer distance to the optimised classification plane indicate a stronger signal of being ISGs or non-ISGs. They are more likely to get higher probability scores (close to 0 or 1) from the SVM model.

**Fig 2.**
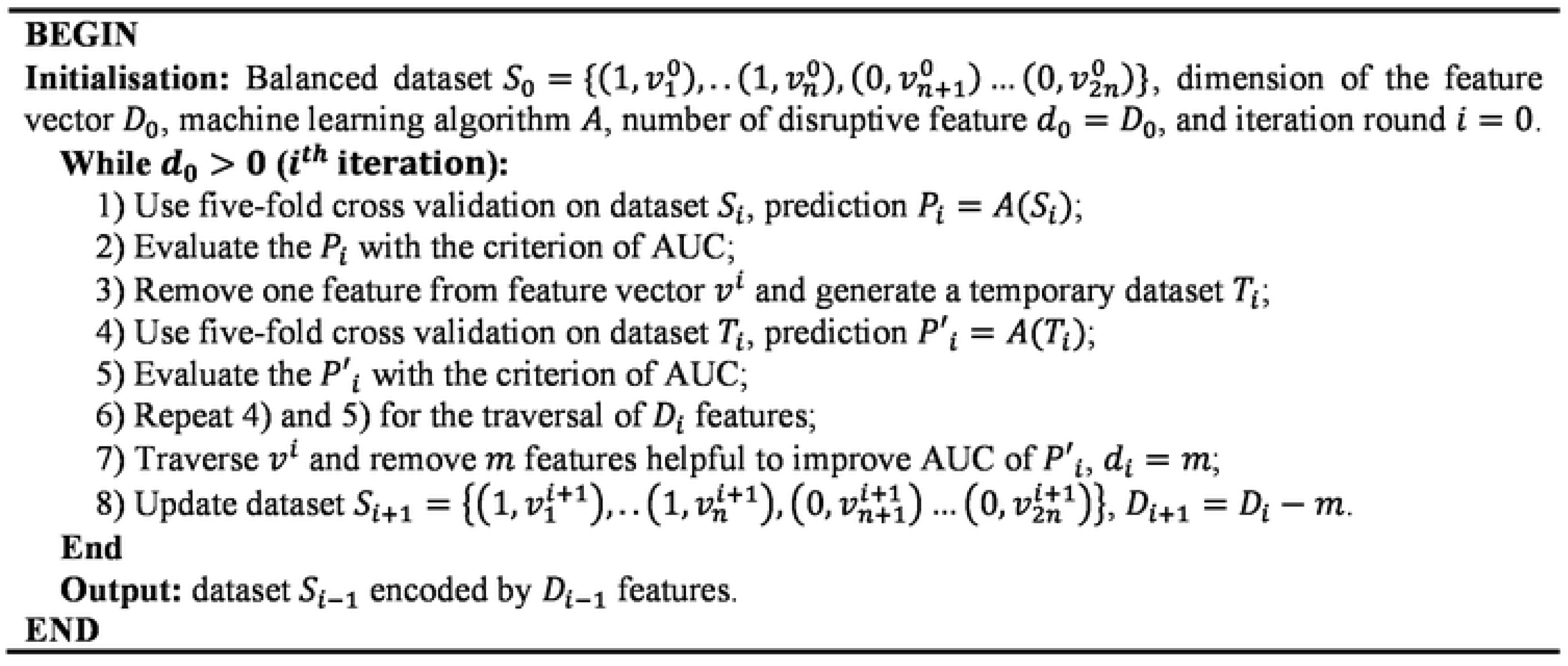
The pseudo-code of the AUC-driven subtractive iteration algorithm. Abbreviations: AUC, area under the receiver operating characteristic curve.

### Performance evaluation

In this study, the prediction results are evaluated with three threshold-dependent criteria, i.e., sensitivity (SN), specificity (SP), and Matthews correlation coefficient (MCC) [54] and two threshold-independent criteria: SN_n and AUC. SN and SP are used to assess the quality of the machine learning model in recognising ISGs and non-ISGs respectively while MCC provides a comprehensive evaluation for both positives and negatives. The number of ‘n’ in the SN_n criterion is determined based on the number of ISGs used for testing. It is used to measure the upper limit of the prediction model as well as to check the existence of important false positives close to the class of ISGs from the perspective of data expression. Finally, AUC is a widely used criterion to evaluate the prediction ability of a binary classifier system. The group of interest is almost unpredictable in a specific binary classifier system if the AUC of the classifier is close to 0.5.

## Results

### Evolutionary characteristics of ISGs

In this study, we construct a dataset consisting of 620 ISGs and 874 non-ISGs (dataset S2) from 10836 well-annotated human genes (dataset S1). Human genes in the S1 dataset have higher confidence based on their records in both the OCISG [8] and Interferome [21] databases. Human genes in both dataset S1 and S2 are evolutionarily unrelated as they are retrieved from the OCISG [8] that compiles clusters of orthologous genes based on whole-genome alignments. However, they may still have inherent characteristics that have resulted in their different expressions in response to the type I IFNs. Here, we explore features relating to polymorphism [34], alternative splicing [35], duplication [37] and mutation [38]. We use the number of ORFs in a human gene to measure its polymorphism. By calculating the average number of ORFs with respect to different Log_2_(Fold Change) levels of expression (window size = 0.1) in the presence of IFNs, we find that human genes with higher Log_2_(Fold Change) tend to have lower levels of polymorphism (Fig 3A). Although low polymorphism seems to be associated with obvious IFN up-regulation, it is not a necessary condition. Compared to the background human genes we include in dataset S1, we find that ISGs tend to have more ORFs, but these differences are not statistically significant (Mann-Whitney U test: *p* > 0.05). We use the number of transcripts to represent the diversity of alternative splicing for a human gene and use the number of protein-coding exons in the canonical transcript to reflect the complexity of the alternative splicing. For these two features, similar negative relationships are observed when Log_2_(Fold Change) increases (Fig 3B & 3C). These results illustrate that the simpler the alternative splicing is, the higher the IFN upregulation. Particularly, as the lowest value of Log_2_(Fold Change) for human genes not differentially expressed only reaches around −0.9. Points placed left of the boundary (x = −0.9) are IRGs. They are generally placed below those with Log_2_(Fold Change) around zero, suggesting the three features (number of ORFs, transcripts and exons) are all differentially represented in some IRGs compared to the remaining non-ISGs. This distribution also indicates that some IRGs have similar feature patterns to ISGs, especially to those highly up-regulated after IFN treatments (right part of the scatter plots in Fig 3).

**Fig 3.**
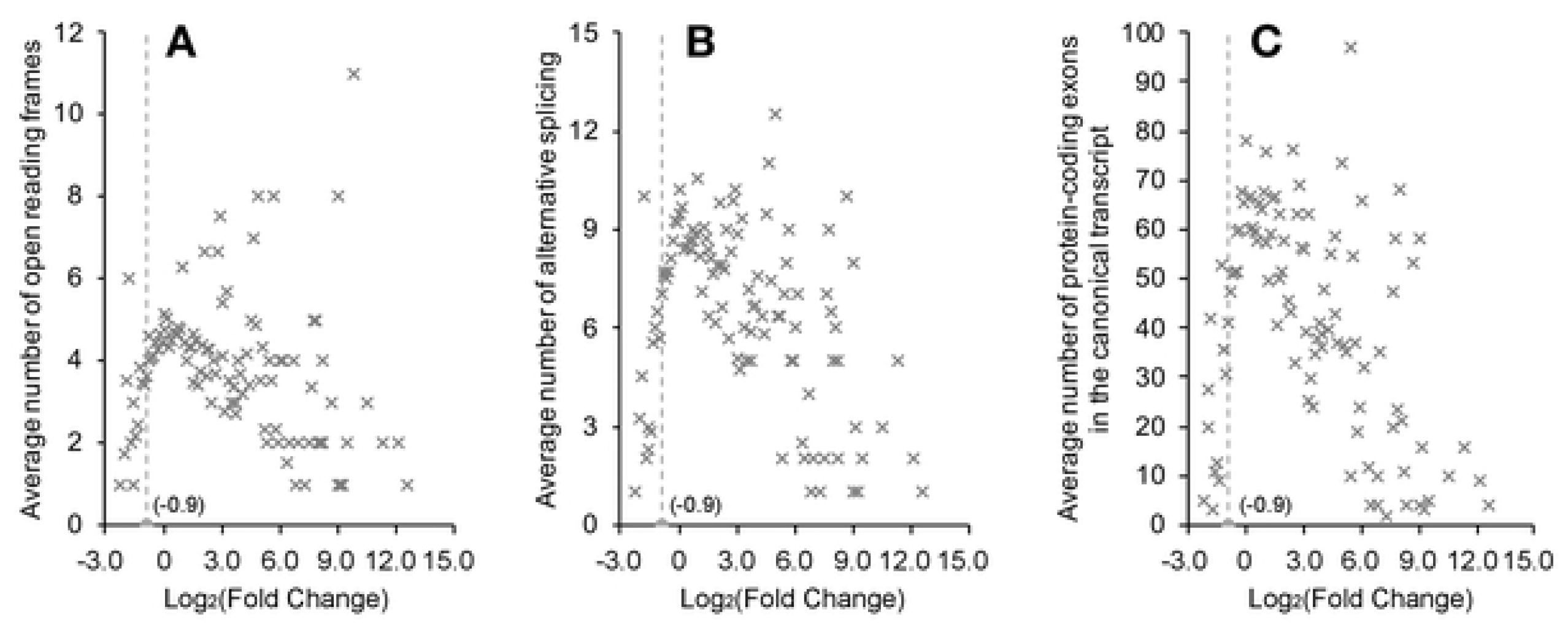
The average representation of features associated with type I IFN stimulations in human fibroblast cells. (A) The number of ORFs as a proxy for gene polymorphism. (B) The number of transcripts as a feature of alternative splicing diversity. (C) The number of protein-coding exons in the canonical transcript as a measurement of the alternative splicing complexity. These three plots are drawn based on the expression data of 8619 human genes with valid fold change in the IFN experiment (**S1 Data**). ELGs are excluded as they have insufficient read coverage to determine a fold change in the IFN experiments. Points in the scatter plot are located based on the average feature representation of genes with similar expression performance in IFN experiments. Abbreviations: IFN, interferon; ORFs, open reading frames; ELGs, human genes with limited expression in interferon experiments.

To determine whether ISGs tend to originate from duplications, we count the number of within human paralogs of each gene (Fig 4A). The results show that there are around 22% of singletons in our main dataset, whilst ISGs have 15% and non-ISGs have 26%. The result of a Mann-Whitney U test [55] indicates that the number of paralogs is significantly under-represented in ISGs compared to the background human genes in dataset S1 (*M_1_* = 10.5, *M_2_* = 11.5, *p* = 8.8E-03). We hypothesize that such a difference is mainly caused by the imbalanced distribution of singletons in ISGs and non-ISGs. The differences become smaller when singletons are excluded from the test (*M_1_* = 12.4, *M_2_* = 14.6, *p* > 0.05). Next, we use the number of non-synonymous substitutions per non-synonymous site (dN) and synonymous substitutions per synonymous site (dS) within human paralogues as a measurement of differences in mutational signatures between different classes [56]. As shown in Fig 4B, non-synonymous substitutions are more frequently observed in ISGs than in background human genes (*M_1_* = 0.62, *M_2_* = 0.55, *p* = 4.0E-03). On the other hand, ISGs also have a higher frequency of synonymous substitutions than background human genes (*M_1_* = 37.7, *M_2_* = 34.6, *p* = 1.1E-02) (Fig 4C) but the difference is not as obvious as for non-synonymous substitutions. The distribution of dN/dS ratios within human paralogues (Fig 4D) indicates that most human genes are constrained by natural selection but ISGs, in general, tend to be less conserved (*M_1_* = 0.036, *M_2_* = 0.045, *p* = 8.3E-03). When eliminating the influence of duplication events, ISGs are still less conserved than non-ISGs but the difference in the dN/dS ratio is not significant (*M_1_* = 0.053, *M_2_* = 0.031, *p* > 0.05).

**Fig 4.**
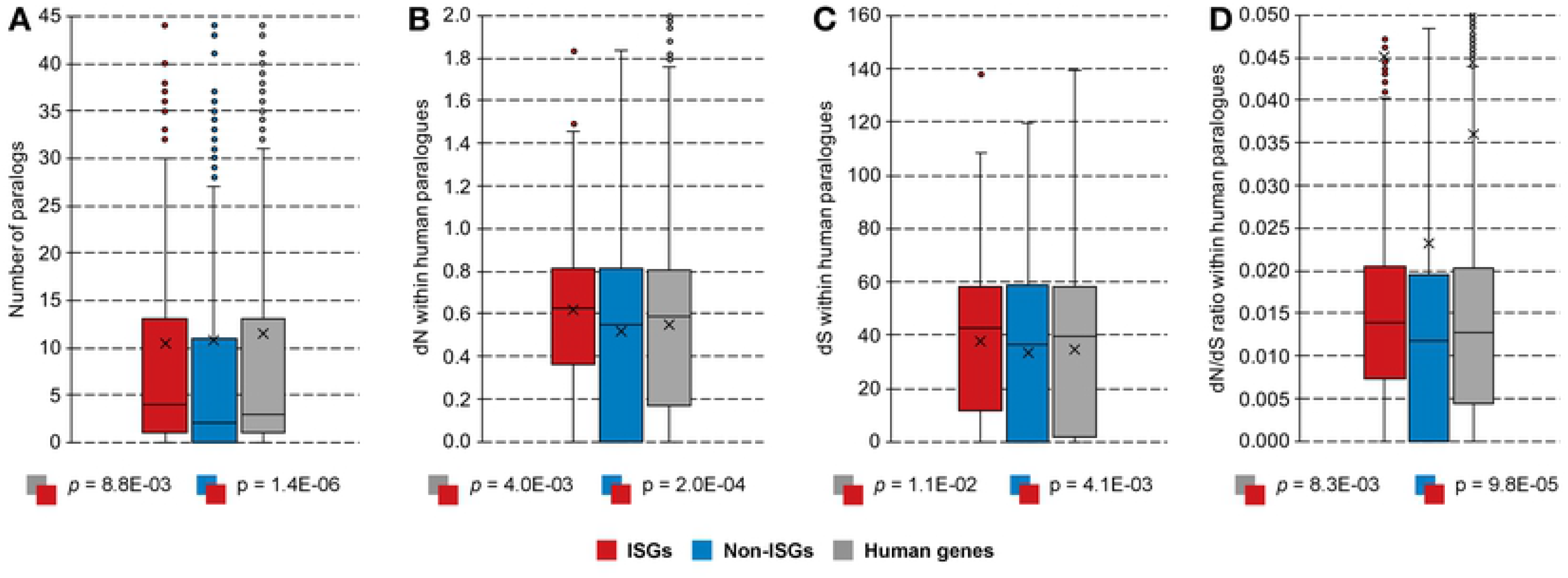
Differences in the evolutionary constraints of human genes. (A) Paralogues within *Homo sapiens*. (B) Non-synonymous substitutions within human paralogues. (C) Synonymous substitutions within human paralogues. (D) dN/dS ratios within human paralogues. Here, ISGs and non-ISGs are taken from dataset S2 while the background human genes are from dataset S1. Mann-Whitney U tests are applied for the hypothesis testing between the feature distribution of different classes. Boxes in the plot represent the major distribution of values (from the first to the third quartile); outliers are added for values higher than two-fold of the third quartile; cross symbol marks the position of the average value including the outliers; upper and lower whiskers show the maximum and minimum values excluding the outliers. Abbreviations: ISGs, interferon upregulated genes; non-ISGs, human genes not significantly up-regulated by interferons; dN, non-synonymous substitutions per non-synonymous site; dS, synonymous substitutions per synonymous site.

### Differences in the coding region of the canonical transcripts

Compared to general profile features (e.g., number of ORFs), the sequences themselves provide more direct mapping to the protein function and structure [57]. Here, we encoded 344 parametric features and 7026 non-parametric features from complementary DNA (cDNA) of the canonical transcript to explore features specific to ISGs. We divide the parametric features into four categories and compare their representations among different classes of human genes, i.e., ISGs, non-ISGs, and the background human genes (Fig 5). Firstly, guanine and cytosine are both more depleted in ISGs than non-ISGs, leading to an under-representation of GC-content in ISGs (Mann-Whitney U test: *M_1_* = 52%, *M_2_* = 55%, *p* = 2.3E-11). This attribute is antithetical to the GC-biased gene conversion (gBGC), making ISGs less stable with weak evolutionary conservation (Fig 4) [58]. Additionally, the under-representation of GC-content also influences the representation of other dinucleotide features. Among all dinucleotide depletions in ISGs, CpG composition is ranked the first followed by GpG and GpC composition (*p* = 2.9E-14, 4.9E-13 and 1.2E-10, respectively). In turn, adenine and thymine-related dinucleotide compositions, exemplified by ApT and TpA are more enriched in ISGs than non-ISGs (*p* = 8.0E-10 and 8.5E-10, respectively).

**Fig 5.**
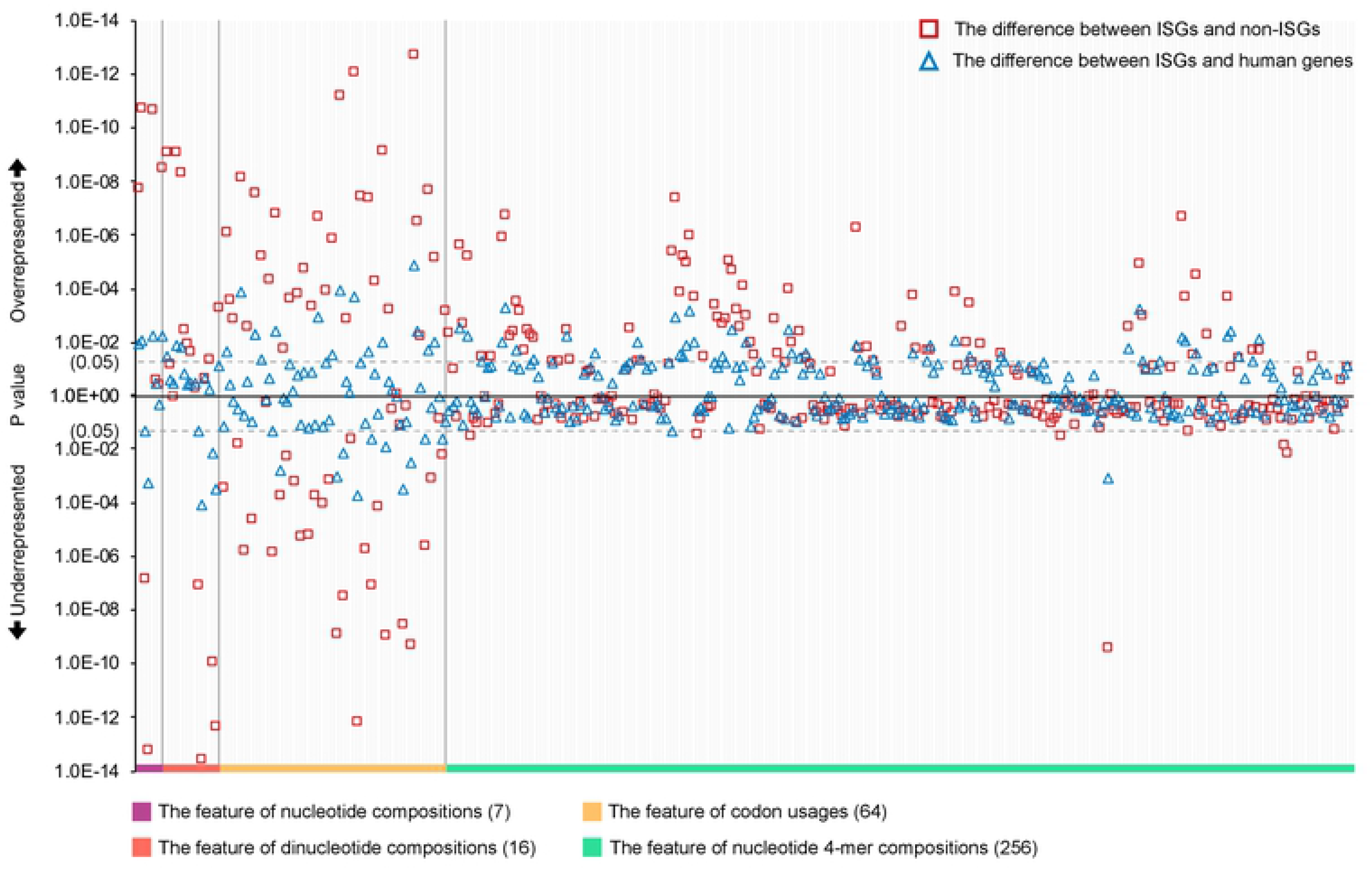
Differences in the representation of parametric features encoded from coding regions of the canonical transcript. Mann-Whitney U tests are applied for hypothesis testing and the results were provided in the **S2 Data**. Here, ISGs and non-ISGs are taken from dataset S2 while the background human genes are from dataset S1. Abbreviations: ISGs, interferon upregulated genes; non-ISGs, human genes not significantly up-regulated by interferons.

Next, we compare the usage of 64 different codons in the third category as their frequencies influence transcription efficiency [43]. Differences between ISGs and background human genes are observed in codons for 11 amino acids including leucine (L), isoleucine (I), valine (V), serine (S), threonine (T), alanine (A), glutamine (Q), lysine (K), glutamic acid (E), arginine (R), and glycine (G). The most significant difference was observed in the usage of codon ‘AGA’. Among all arginine-targeted alternative codons, codon ‘AGA’ is usually favoured, and its usage reaches an estimated 25% in ISGs, but reduces to 22% in the background human genes and is even significantly lower in non-ISGs, at 18% (*p* = 1.4E-05 and 1.9E-13, respectively). On the other hand, compared to background human genes, the codon ‘CAG’ coding for amino acid ‘Q’ is the most under-represented in ISGs. It is less favoured by ISGs than non-ISGs (*M_1_* = 72%, *M_2_* = 78%, *p* = 7.3E-13) although it dominates in coding patterns. As for the three stop codons, comparing with background human genes, the usage of the ochre stop codon, i.e., ‘TAA’ is over-represented in ISGs (*M_1_* = 28%, *M_2_* = 33%, *p* = 9.7E-03). In this category of codon usage, the features that have different frequencies between ISGs and background human genes became more discriminating when comparing ISGs with non-ISGs. Significant differences in codon usages between ISGs and non-ISGs are widely observed except for methionine (M) and tryptophan (W). Hence, although we find a limited number of codon usage features showing significant differences between ISGs and the background human genes, the codon usage features are useful for discriminating ISGs from non-ISGs.

In the last category, we calculate the occurrence frequency of 256 nucleotide 4-mers to add some positional resolution for finding and comparing interesting organisational structures [41]. Among the 256 4-mers, we find 46 of them are differentially represented between ISGs and background human genes (**S2 Data**). Most of these 4-mers are over-represented by ISGs except two with the pattern ‘TAAA’ and ‘CGCG’. Interestingly, the feature of ‘TAAA’ composition becomes a positive factor when comparing ISGs and non-ISGs (*M_1_* = 4.1%, *M_2_* = 3.7%, *p* = 4.1E-06), suggesting it may be a good feature to ascertain potential or incorrectly labelled ISGs. We find six nucleotide 4-mers: ‘ACCC’, ‘AGTC’, ‘AGTG’, ‘TGCT’, ‘GACC’, and ‘GTGC’ are over-represented in ISGs when compared to background human genes but are not differentially represented when comparing ISGs with non-ISGs. Thus, these six features may be inherently biased for some unknown reasons and are not powerful enough to distinguish ISGs from non-ISGs. In addition to the aforementioned 40 features that are over-represented in ISGs compared to background human genes, we find a further 39 features nucleotide 4-mers differentially represented between ISGs and non-ISGs (**S2 Data**).

To check the effect of these aforementioned 343 features on the level of stimulation in the IFN system (Log_2_(Fold Change) > 0), we calculate the PCC for the normalised features (**Equation 2**) and find 106 features are positively related to the increase of fold change, and 34 features are suppressed when human gene are more up-regulated (Student t-test: *p* < 0.05) (**S3 Data**). ApA composition shows the most obvious positive correlation with stimulation level (PCC = 0.464, *p* = 8.8E-06) while negative association between the representation of 4-mer ‘CGCG’ and IFN-induced up-regulation is the most significant (PCC = −0.593, *p* = 3.2E-09). Human genes with higher up-regulation in the presence of IFNs contain more codons ‘CAA’ rather than ‘CAG’ for coding amino acid ‘Q’. The depletion of GC-content, especially cytosine content, promotes the suppression of many nucleotide compositions in the cDNA, e.g. CpG composition.

To find conserved sequence patterns related to gene regulations [59], we check the existence of 2940, 44100 and 661500 short linear nucleotide motifs (SLim_DNAs) consisting of three to five consecutive nucleobases in the group of ISGs and non-ISGs. By using a positive 5% difference in the occurrence frequency as cut-off threshold, we find 7884 SLim_DNAs with a maximum difference in representation around 15%. After using Pearson’s chi-squared tests and Benjamini-Hochberg correction to avoid type I error in multiple hypotheses [52], 7025 SLim_DNAs remain with an adjusted p-value lower than 0.01 (**S4 Data**), hereon referred to as flagged SLim_DNAs. Here, the differentially represented 7025 SLim_DNAs are ranked according to the adjusted p-value. As shown in Fig 6A, dinucleotide ‘TpA’ dominates in the top 10, top 100, top 1000, and all differentially represented SLim_DNAs even if TpA representation is suppressed in the cDNA of genes’ canonical transcripts compared to other dinucleotides. Dinucleotide ‘ApT’ and ‘ApA’ are also frequently observed in the flagged SLim_DNAs but their occurrences do not show significant difference in the top 100 SLim_DNAs (Pearson’s chi-squared test: *p* > 0.05). GC-related dinucleotides, e.g., ‘CpC’, ‘GpC’ and ‘GpG’ are rarely observed in the flagged SLim_DNAs especially in the top 10 or top 100. In view of these, we hypothesize that the differential representation of nucleotide compositions influences and reflects on the pattern of SLim_DNAs in ISGs. By checking the co-occurrence status of the flagged SLim_DNAs, we find these sequence patterns have a cumulative effect in distinguishing ISGs from non-ISGs especially when the number of cooccurring SLim_DNAs reaches around 5320 (Pearson’s chi-squared test: *p* = 7.9E-13, Fig 6B). There are eight (∼1.3%) ISGs in the refined set, i.e., dataset S2 containing all the flagged 7025 SLim_DNAs. Their up-regulation after IFN treatment are generally low with a fold change fluctuating around 2.2. Although some genes such as desmoplakin (DSP) are clearly highly up-regulated in endothelial cells isolated from human umbilical cord veins after both IFN-α (fold change = 11.1) and IFN-β (fold change = 13.7) treatments. We also find some non-ISGs and ELGs containing the flagged SLim_DNAs, e.g., hemicentin 1 (HMCN1) and tudor domain containing 6 (TDRD6), but their frequencies are lower than that in ISGs. Although there is an obvious imbalance between the number of ISGs and non-ISGs in the human genome [9–11], the curve for the background human genes in Fig 6B is still closer to that for ISGs rather than that for non-ISGs. It suggests that some genetic patterns are widely represented in the coding region of human genes, making them potentially up-regulated in the IFN system.

**Fig 6.**
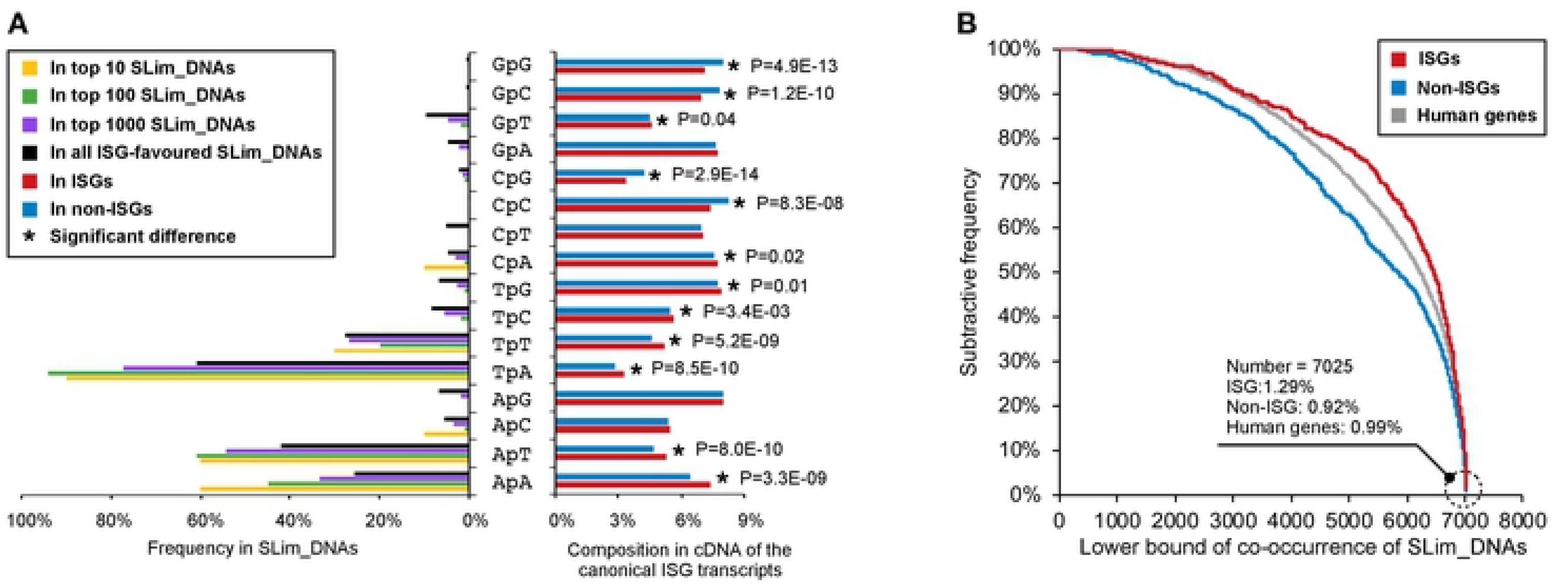
The pattern of SLim_DNAs in the coding region of the canonical transcripts. (A) Influence of dinucleotide compositions on the flagged SLim_DNAs. (B) The co-occurrence status of SLim_DNAs in different human genes. Ranks in (A) are generated based on the adjust p value given by Pearson’s chi-squared tests after Benjamini-Hochberg correction procedure. Detailed results of the hypothesis tests are provided in **S4 Data**. Here, ISGs and non-ISGs are taken from dataset S2 while the background human genes are from dataset S1. Abbreviations: ISGs, interferon-stimulated genes; non-ISGs, human genes not significantly up-regulated by interferons; SLim_DNAs, short linear nucleotide motifs; cDNA, complementary DNA.

### Differences in the protein sequence

We use the protein sequences generated by the canonical transcript to extract features at the proteomic level. In addition to the basic composition of 20 standard amino acids, we consider 17 additional features related to physicochemical (e.g., hydropathy and polarity) or geometric properties (e.g., volume) [60, 61]. We find several amino acids that are either enriched or depleted in ISG products compared to background human proteins, which are produced by genes in dataset S1 (Fig 7). The differences are even more marked between protein products of ISGs and non-ISGs, highlighting some differences that are not observed when comparing ISG products to the background human proteins (e.g., isoleucine composition). The differences observed in the amino acid compositions are at least in part associated with the patterns previously observed in features encoded from genetic coding regions. For example, asparagine (N) shows significant over-representation in ISG products compared to non-ISG products or background human proteins (Mann-Whitney U test: *p* = 2.8E-12 and 1.2E-03, respectively). This is expected as there are only two codons, i.e., ‘AAT’ and ‘AAC’ coding for amino acid ‘N’, and dinucleotide ‘ApA’ shows a remarkable enrichment in the coding region of ISGs. A similar explanation can be given for the relationship between the deficiency of GpG content and amino acid ‘G’. The translation of amino acid ‘K’ is also influenced by ApA composition but is not significant due to the mild representation of dinucleotide ‘ApG’ in the genetic coding region. Additionally, as previously mentioned, ISGs show a significant depletion in the CpG content, and consequently, the amino acid ‘A’ and ‘R’ in ISG products are significantly under-represented. Cysteine (C) is not frequently observed in human proteins but still shows a relatively significant enrichment in ISG products (*M_1_* = 2.3%, *M_2_* = 2.5%, *p* = 1.8E-03).

**Fig 7.**
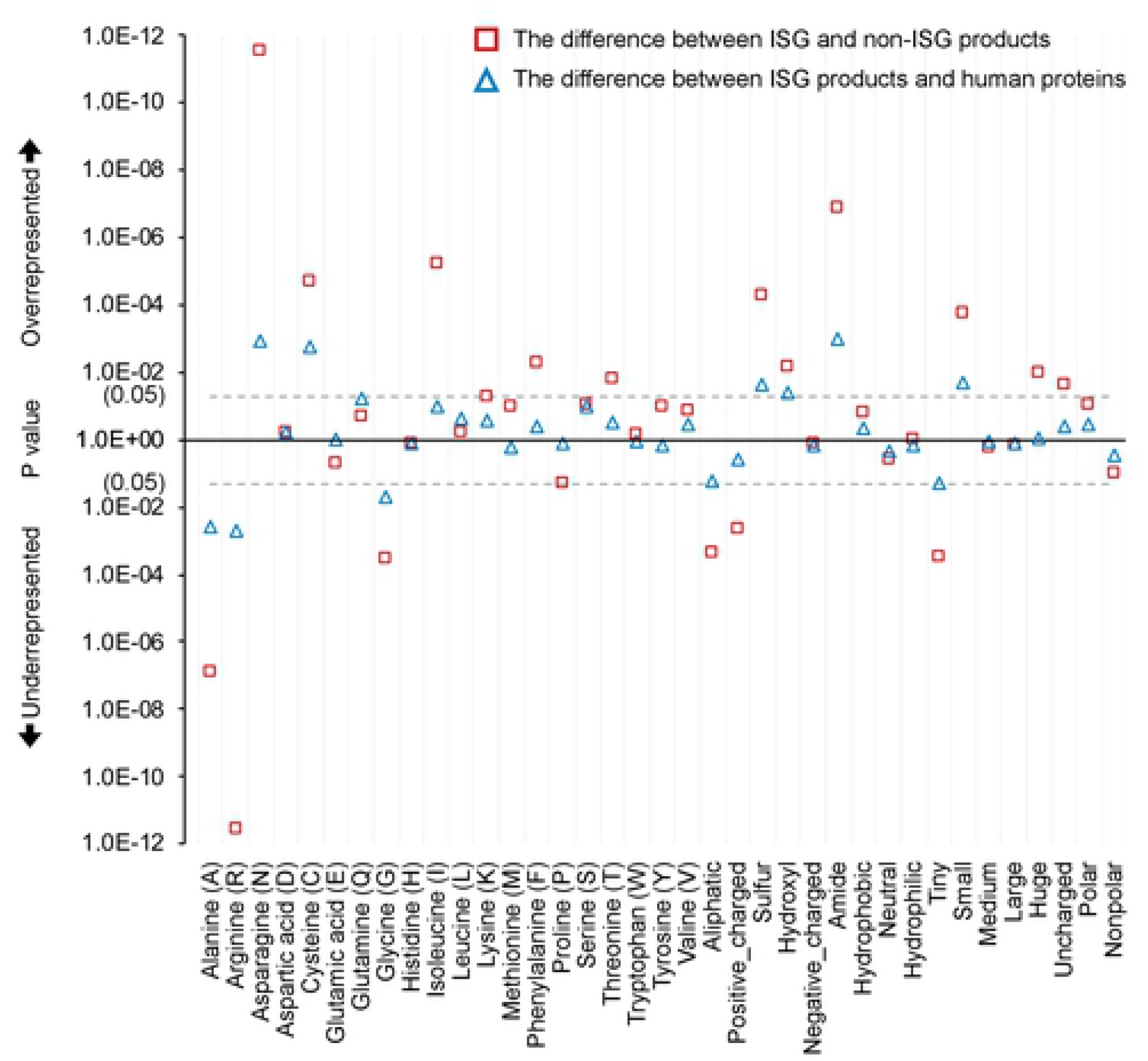
Differences in the representation of parametric features encoded from protein sequences. Mann-Whitney U tests are applied for hypothesis testing and the results were provided in the **S2 Data**. Here, ISGs and non-ISGs are taken from dataset S2 while the background human genes are from dataset S1. Aliphatic group: amino acid ‘A’, ‘G’, ‘I’, ‘L’, ‘P’ and ‘V’; aromatic/huge group: amino acid ‘F’, ‘W’ and ‘Y’ (volume > 180 cubic angstroms); sulfur group: amino acid ‘C’ and ‘M’; hydroxyl group: amino acid ‘S’ and ‘T’; acidic/negative_charged group: amino acid ‘D’ and ‘E’; amide group: amino acid ‘N’ and ‘Q’; positive_charged group: amino acid ‘R’, ‘H’ and ‘K’; hydrophobic group: amino acid ‘A’, ‘C’, ‘I’, ‘L’, ‘M’, ‘F’, ‘V’, and ‘W’ that participates to the hydrophobic core of the structural domains [44]; neutral group: amino acid ‘G’, ‘H’, ‘P’, ‘S’, ‘T’ and ‘Y’; hydrophilic group: amino acid ‘R’, ‘N’, ‘D’, ‘Q’, ‘E’ and ‘K’; Tiny group: amino acid ‘G’, ‘A’ and ‘S’ (volume < 90 cubic angstroms); small group: amino acid ‘N’, ‘D’, ‘C’, ‘P’ and ‘T’ (volume ranged from 109 to 116 cubic angstroms); medium group: amino acid ‘Q’, ‘E’, ‘H’ and ‘V’ (volume ranged within 138 to 153 cubic angstroms); large group: amino acid ‘R’, ‘I’, ‘L’, ‘K’ and ‘M’ (volume ranged within 163 to 173 cubic angstroms); uncharged group: the remaining 15 amino acids except electrically charged ones; polar group: amino acid ‘R’, ‘H’, ‘K’, ‘D’, ‘E’, ‘N’, ‘Q’, ‘S’, ‘T’ and ‘Y’; nonpolar group: the remaining 10 amino acids except polar ones. Abbreviations: ISG, interferon upregulated genes; non-ISG, human genes not significantly up-regulated by interferons.

When focusing on the composition of amino acids grouped by physicochemical or geometric properties, we also find some features differentially represented between ISG products and background human proteins. The result shows that hydroxyl (amino acid ‘S’ and ‘T’), amide (amino acid ‘N’ and ‘Q’), or sulfur amino acids (amino acid ‘C’ and ‘M’) are more abundant in ISG products compared to the background human proteins (Mann-Whitney U test: *p* = 0.04, 1.0E-03 and 0.02, respectively). Small amino acids (amino acid ‘N’, ‘C’, ‘T’, aspartic acid (D) and proline (P), the volume ranges from 108.5 to 116.1 cubic angstroms) are more frequently observed in ISG products than in background human proteins (*M_1_* = 22.1%, *M_2_* = 21.7%, *p* = 0.02). The differences become more marked when comparing the representation of these features between ISG and non-ISG products. For example, features relating to chemical properties of the side chain (e.g., aliphatic), charge status and geometric volume show differences between proteins produced by ISGs and non-ISGs. Some features, e.g., neutral amino acids that include amino acid ‘G’, ‘P’, ‘S’, ‘T’, histidine (H) and tyrosine (Y) are not differentially represented between ISG and non-ISG products, but they show obvious association with the change of IFN-triggered stimulations (PCC = −0.556, *p* = 4.1E-08) (**S3 Data**).

We then search the sequence of ISG products against that of non-ISG products to find conserved short linear amino acid motifs (SLim_AAs), which may have resulted from strong purifying selection [45]. As opposed to the analysis on the genetic sequence, we only obtain 19 enriched sequence patterns with a Pearson’s chi-squared p value ranging from 1.5E-04 to 0.02 (Table 2). These SLim_AAs are greatly influenced by four polar amino acids: ‘K’, ‘N’, ‘E’ and ‘S’, and one nonpolar amino acid: ‘L’. Some of these SLim_AAs, e.g., SLim ‘NVT’ and ‘S-N-E’, are clearly over-represented in ISG products compared to background human proteins and can be used as features to differentiate ISGs from background human genes. The third column in Table 2 also indicates a number of patterns, e.g., SLim ‘S-N-T’, that are lacking in non-ISG products and hence may be the reason for the lack of up-regulation in the presence of IFNs. Particularly, we noticed that SLim ‘KEN’ is a destruction motif that can be recognised or targeted by anaphase promoting complex (APC) for polyubiquitination and proteasome-mediated degradation [62, 63]. Results shown in Fig 8A illustrate that the co-occurrence of differentially represented SLim_AAs has a cumulative effect in distinguishing ISGs from non-ISGs. This cumulative effect can be achieved with only two random SLim_AAs (Pearson’s chi-squared test: *p* = 4.6E-10). The bias in the co-occurring SLim_AAs in the background human proteins towards a pattern similar to non-ISG products further proves the importance of these 19 SLim_AAs. However, their co-occurrence is not associated with the level of IFN-triggered stimulations (PCC = 0.015, *p* > 0.05) (Fig 8B)

**Fig 8.**
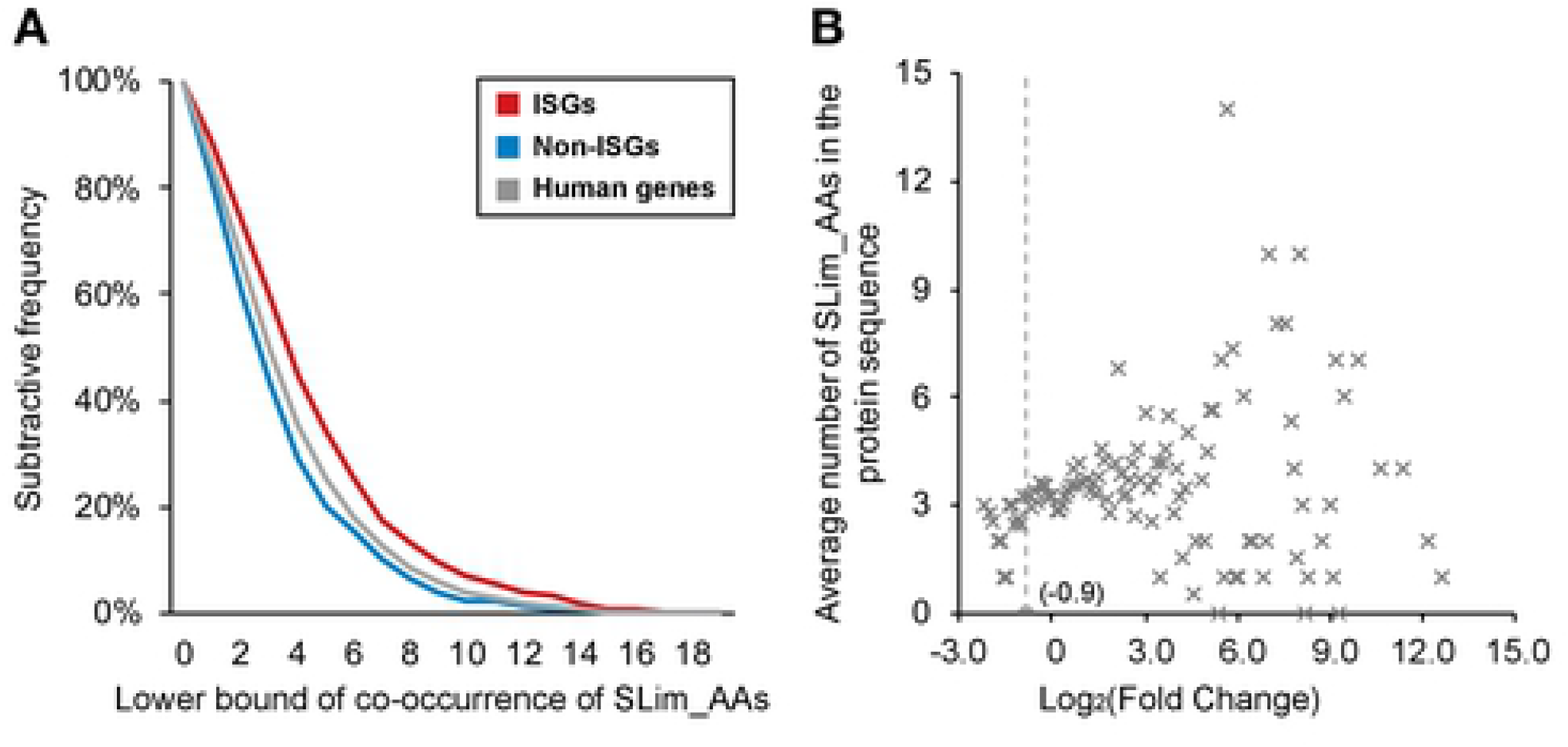
Representation of co-occurred SLim_AAs in our main dataset. (A) The co-occurrence status of SLim_AAs in different classes. (B) Relationship between co-occurrence of the marked SLim_AAs and Log_2_(Fold Change) after IFN treatments. Here, ISGs and non-ISGs are taken from dataset S2 while the background human genes are from dataset S1. Points in (B) are located based on the average feature representation of genes with similar expression performance in IFN experiments. Abbreviations: IFN, interferon; ISGs, interferon-stimulated genes; non-ISGs, human genes not significantly up-regulated by interferons; SLim_AAs, short linear amino acid motifs.

**Table 2.**
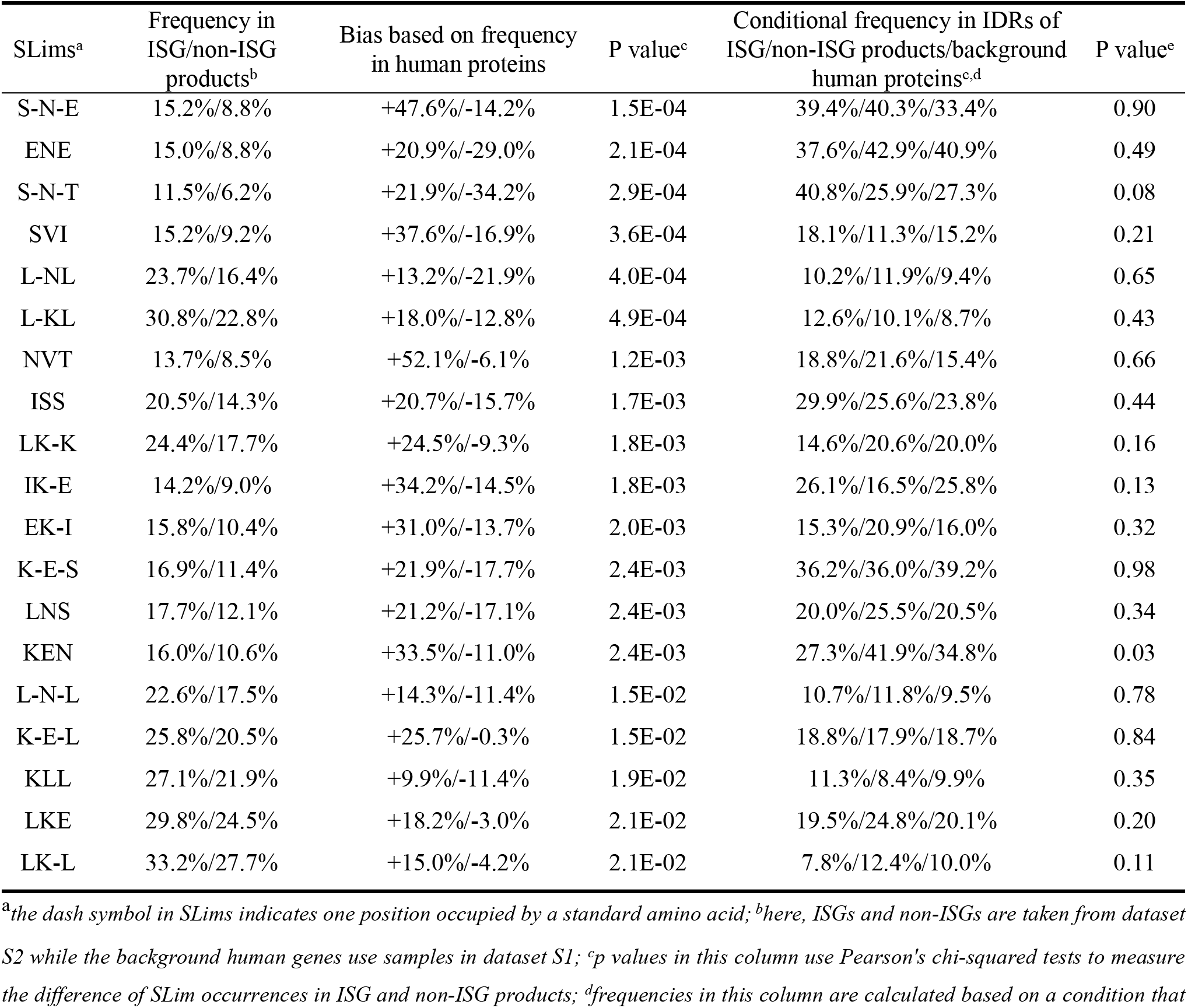
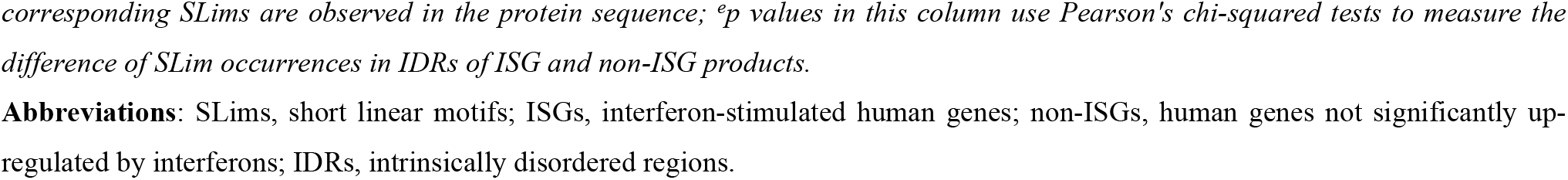
Representation of SLims in protein sequences and their IDRs.

| SLims<sup>a</sup> | Frequency in ISG/non-ISG products<sup>b</sup> | Bias based on frequency in human proteins | P value<sup>c</sup> | Conditional frequency in IDRs of ISG/non-ISG products/background human proteins<sup>c,d</sup> | P value<sup>e</sup> |
| --- | --- | --- | --- | --- | --- |
| S-N-E | 15.2%/8.8% | +47.6%/-14.2% | 1.5E-04 | 39.4%/40.3%/33.4% | 0.90 |
| ENE | 15.0%/8.8% | +20.9%/-29.0% | 2.1E-04 | 37.6%/42.9%/40.9% | 0.49 |
| S-N-T | 11.5%/6.2% | +21.9%/-34.2% | 2.9E-04 | 40.8%/25.9%/27.3% | 0.08 |
| SVI | 15.2%/9.2% | +37.6%/-16.9% | 3.6E-04 | 18.1%/11.3%/15.2% | 0.21 |
| L-NL | 23.7%/16.4% | +13.2%/-21.9% | 4.0E-04 | 10.2%/11.9%/9.4% | 0.65 |
| L-KL | 30.8%/22.8% | +18.0%/-12.8% | 4.9E-04 | 12.6%/10.1%/8.7% | 0.43 |
| NVT | 13.7%/8.5% | +52.1%/-6.1% | 1.2E-03 | 18.8%/21.6%/15.4% | 0.66 |
| ISS | 20.5%/14.3% | +20.7%/-15.7% | 1.7E-03 | 29.9%/25.6%/23.8% | 0.44 |
| LK-K | 24.4%/17.7% | +24.5%/-9.3% | 1.8E-03 | 14.6%/20.6%/20.0% | 0.16 |
| IK-E | 14.2%/9.0% | +34.2%/-14.5% | 1.8E-03 | 26.1%/16.5%/25.8% | 0.13 |
| EK-I | 15.8%/10.4% | +31.0%/-13.7% | 2.0E-03 | 15.3%/20.9%/16.0% | 0.32 |
| K-E-S | 16.9%/11.4% | +21.9%/-17.7% | 2.4E-03 | 36.2%/36.0%/39.2% | 0.98 |
| LNS | 17.7%/12.1% | +21.2%/-17.1% | 2.4E-03 | 20.0%/25.5%/20.5% | 0.34 |
| KEN | 16.0%/10.6% | +33.5%/-11.0% | 2.4E-03 | 27.3%/41.9%/34.8% | 0.03 |
| L-N-L | 22.6%/17.5% | +14.3%/-11.4% | 1.5E-02 | 10.7%/11.8%/9.5% | 0.78 |
| K-E-L | 25.8%/20.5% | +25.7%/-0.3% | 1.5E-02 | 18.8%/17.9%/18.7% | 0.84 |
| KLL | 27.1%/21.9% | +9.9%/-11.4% | 1.9E-02 | 11.3%/8.4%/9.9% | 0.35 |
| LKE | 29.8%/24.5% | +18.2%/-3.0% | 2.1E-02 | 19.5%/24.8%/20.1% | 0.20 |
| LK-L | 33.2%/27.7% | +15.0%/-4.2% | 2.1E-02 | 7.8%/12.4%/10.0% | 0.11 |

<sup>a</sup>the dash symbol in SLims indicates one position occupied by a standard amino acid; <sup>b</sup>here, ISGs and non-ISGs are taken from dataset S2 while the background human genes use samples in dataset S1; <sup>c</sup>p values in this column use Pearson's chi-squared tests to measure the difference of SLim occurrences in ISG and non-ISG products; <sup>d</sup>frequencies in this column are calculated based on a condition that
corresponding SLims are observed in the protein sequence; <sup>e</sup>p values in this column use Pearson's chi-squared tests to measure the difference of SLim occurrences in IDRs of ISG and non-ISG products. **Abbreviations:** SLims, short linear motifs; ISGs, interferon-stimulated human genes; non-ISGs, human genes not significantly upregulated by interferons; IDRs, intrinsically disordered regions.

Regions that lacked stable structures under normal physiological conditions within proteins are termed intrinsically disordered regions (IDRs). They play an important role in cell signalling [64]. Compared with ordered regions, IDRs are usually more accessible and have multiple binding motifs, which can potentially bind to multiple partners [65]. According to the results calculated by IUPred [66], we find 6721, 10510, and 119071 IDRs (IUpred score no less than 0.5) in proteins produced by ISGs, non-ISGs and background human genes respectively. We hypothesize that enriched SLims widely detected in IDRs may be important for human protein-protein interactions or potentially virus mimicry [51]. For instance, in ISG products, 29 out of 71 SLim ‘S-N-T’ are observed in IDRs (∼40.8%), 14.9% higher than that in non-ISG products (Table 2). This difference reflects the importance of SLim ‘S-N-T’ for target specificity of IFN-induced protein-protein interactions [9] even if it is not statistically significant. By contrast, the conditional frequency of SLim ‘S-N-E’ is discovered in IDRs of ISG and non-ISG products are almost the same, indicating that SLim ‘S-N-E’ may have an association with some inherent attributes of ISGs but is less likely to be involved in IFN-induced protein-protein interactions. SLim ‘KEN’ in IDRs also shows some interesting differences: in non-ISG products, 41.9% of SLim ‘KEN’ are observed in IDRs, 14.6% higher than that in ISG products, which provides an effective approach to distinguish ISGs from non-ISGs. When SLim ‘KEN’ is discovered in the ordered region of a protein sequence, statistically, the protein is more likely to be produced by an ISG, but this assumption is reversed if the SLim is located in an IDR (Pearson’s chi-squared tests: *p* = 0.03). Despite the relatively low conditional frequency of SLim ‘KEN’ in the IDRs of ISG products, these SLim_AAs in the IDR are more likely to be functionally active than those falling within ordered globular regions [67].

### Differences in network profiles

We construct a network with 332,698 experimentally verified interactions among 17603 human proteins (confidence score > 0.63) from the HIPPIE database [47]. 10169 out of 10836 human proteins from our background dataset S1 are included in it. Nodes and edges of this network can be downloaded from our webserver at http://isgpre.cvr.gla.ac.uk/. Based on this network, we calculate eight features including the average shortest path, closeness, betweenness, stress, degree, neighbourhood connectivity, clustering coefficient, and topological coefficient. As illustrated in Fig 9, ISG products tend to have higher values of betweenness and stress than background human proteins (Mann-Whitney U test: *p* = 0.01, and 0.03, respectively), which means they are more likely to locate at key paths connecting different nodes of the PPI network. Some ISG products with high values of betweenness and stress, e.g., tripartite motif containing 25 (TRIM25), can be considered as the shortcut or bottleneck of the network and play important roles in many PPIs including those related to the IFN-triggered immune activities [68, 69]. However, the over-representation of betweenness does not mean ISG products are more likely to be or even be close to bottlenecks in the network compared to background human proteins. Some examples shown in Table 3 indicate that ISG products are less-connected by top-ranked bottlenecks and hubs in the network than non-ISGs or background human proteins. This conclusion is not influenced by hub/bottleneck protein’s performance in the IFN experiments. Comparing proteins produced by ISGs and non-ISGs, we find the former tends to have lower values of clustering coefficient and neighbourhood connectivity (Mann-Whitney U test: *p* = 0.04, and 7.9E-03, respectively), which means that ISG products and the majority of their interacting proteins are less likely to be targeted by lots of proteins. It also supports the finding that ISG products are involved in many shortest paths for nodes but are away from hubs or bottlenecks in the network. To some extents, this location also increases the length of the average shortest paths through ISG products in the network.

**Fig 9.**
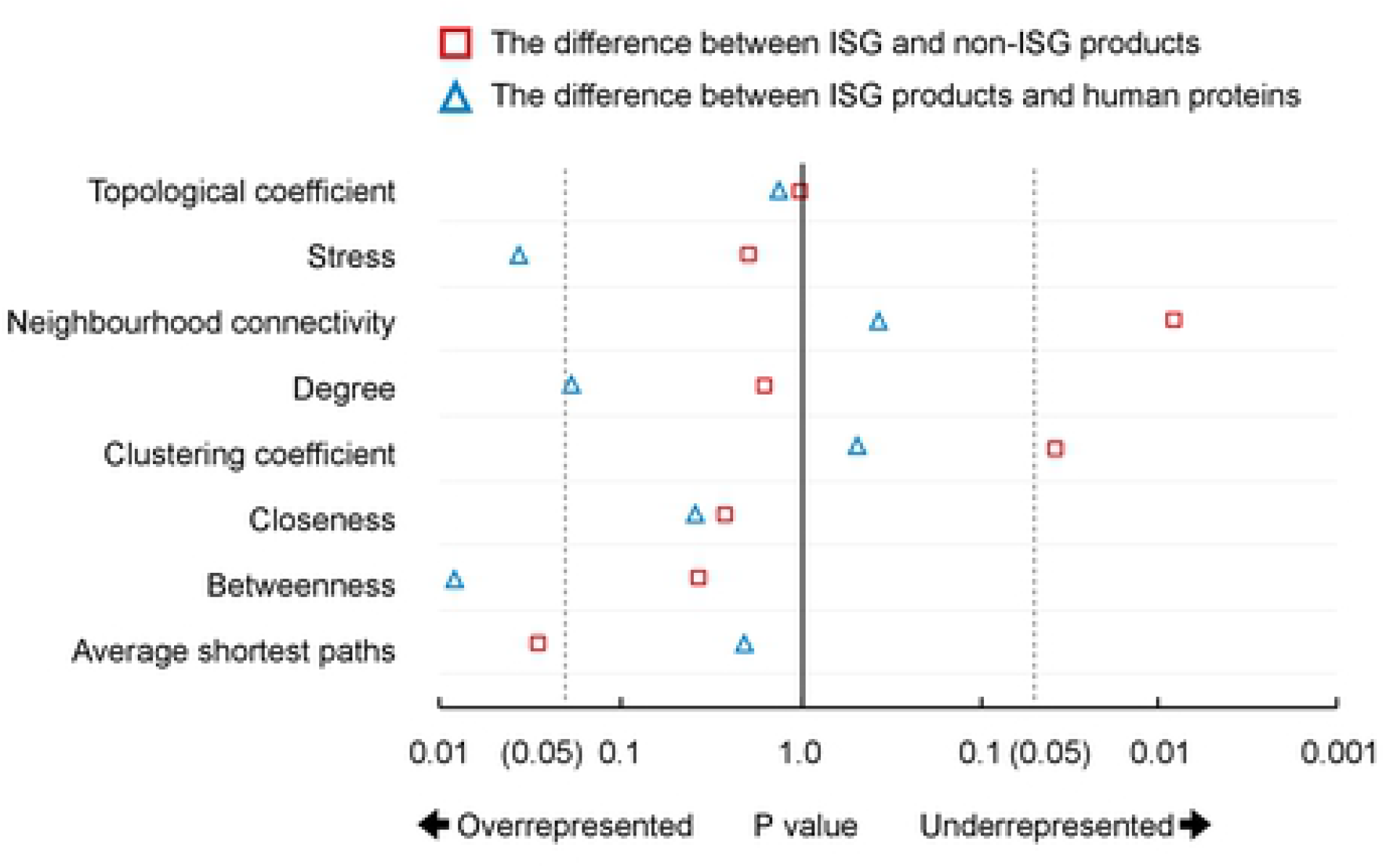
Differential network preferences of proteins coded by different human genes. Mann-Whitney U tests are applied for hypothesis testing and the results were provided in the **S2 Data**. Here, ISGs and non-ISGs are taken from dataset S2 while the background human genes use samples in dataset S1. Abbreviations: ISGs, interferon-stimulated genes; non-ISGs, human genes not significantly up-regulated by interferons.

**Table 3.**
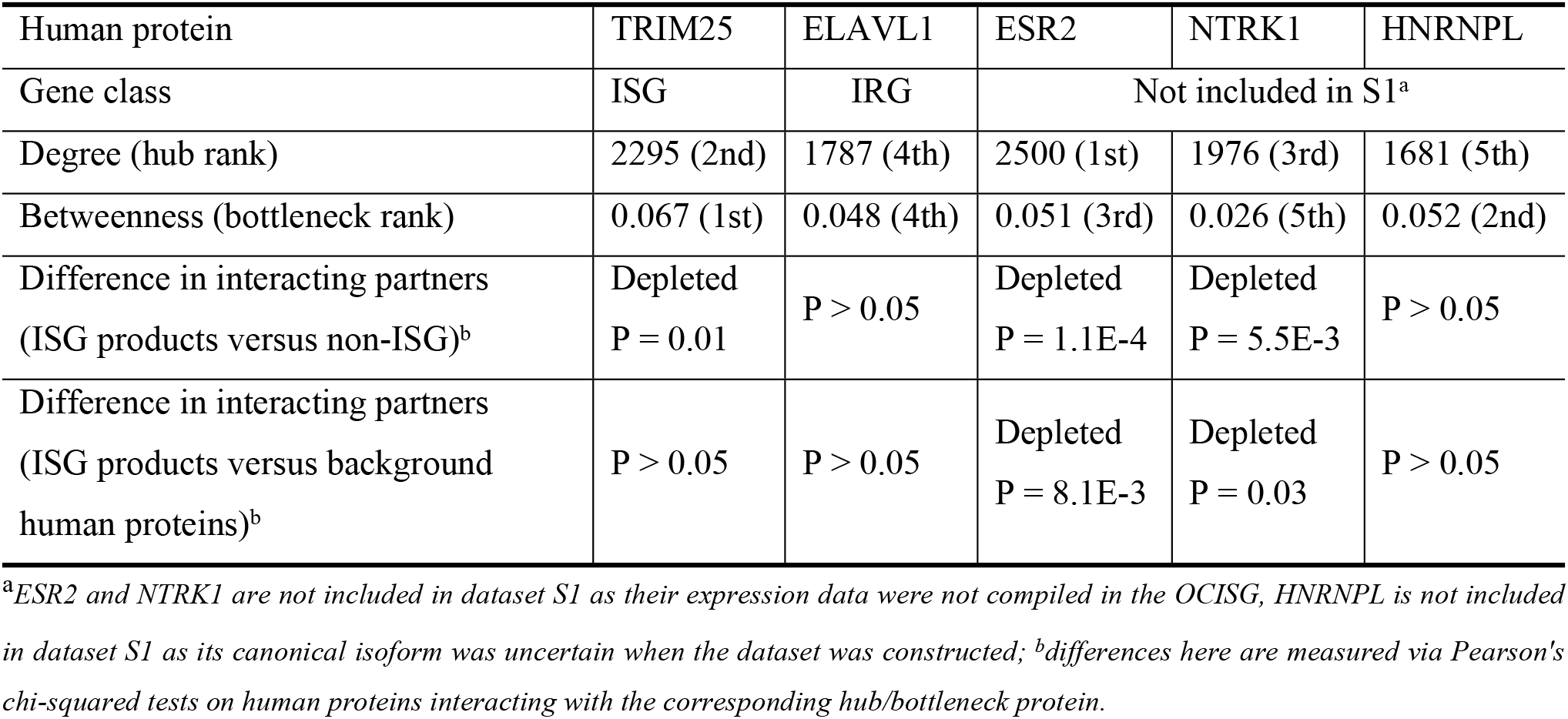
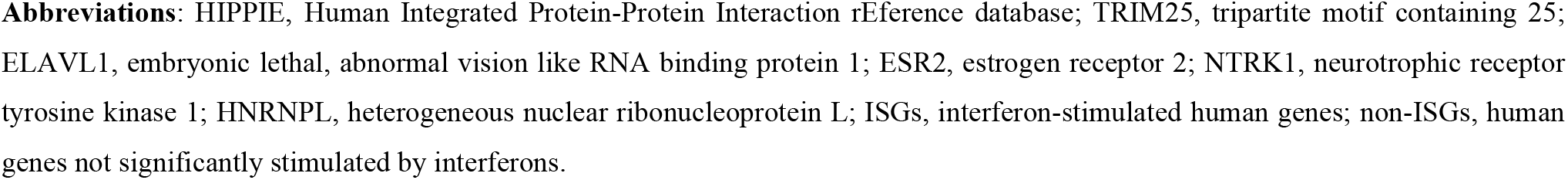
Interaction profiles of human proteins connecting top hubs/bottlenecks of the HIPPIE network.

| Human protein | TRIM25 | ELAVL1 | ESR2 | NTRK1 | HNRNPL |
| --- | --- | --- | --- | --- | --- |
| Gene class | ISG | IRG | Not included in S1<sup>a</sup> |  |  |
| Degree (hub rank) | 2295 (2nd) | 1787 (4th) | 2500 (1st) | 1976 (3rd) | 1681 (5th) |
| Betweenness (bottleneck rank) | 0.067 (1st) | 0.048 (4th) | 0.051 (3rd) | 0.026 (5th) | 0.052 (2nd) |
| Difference in interacting partners (ISG products versus non-ISG)<sup>b</sup> | Depleted<br>P = 0.01 | P > 0.05 | Depleted<br>P = 1.1E-4 | Depleted<br>P = 5.5E-3 | P > 0.05 |
| Difference in interacting partners (ISG products versus background human proteins)<sup>b</sup> | P > 0.05 | P > 0.05 | Depleted<br>P = 8.1E-3 | Depleted<br>P = 0.03 | P > 0.05 |

<sup>a</sup>ESR2 and NTRK1 are not included in dataset S1 as their expression data were not compiled in the OCISG, HNRNPL is not included in dataset S1 as its canonical isoform was uncertain when the dataset was constructed; <sup>b</sup>differences here are measured via Pearson's chi-squared tests on human proteins interacting with the corresponding hub/bottleneck protein.
**Abbreviations:** HIPPIE, Human Integrated Protein-Protein Interaction rEference database; TRIM25, tripartite motif containing 25; ELAVL1, embryonic lethal, abnormal vision like RNA binding protein 1; ESR2, estrogen receptor 2; NTRK1, neurotrophic receptor tyrosine kinase 1; HNRNPL, heterogeneous nuclear ribonucleoprotein L; ISGs, interferon-stimulated human genes; non-ISGs, human genes not significantly stimulated by interferons.

When investigating the association between IFN-induced gene stimulation and network attributes of gene products, we only find the feature of neighbourhood connectivity is under-represented as the level of differential expression in the presence of IFN increases (PCC = −0.392, *p* = 2.2E-04). This suggests that proteins produced by genes that are highly up-regulated in response to IFNs are further away from hubs in the PPI networks.

### Features highly associated with the level of IFN stimulations

In this study, we encode a total of 397 parametric and 7046 non-parametric features covering the aspects of evolutionary conservation, nucleotide composition, transcription, amino acid composition, and network profiles. In order to find out some key factors that may enhance or suppress the stimulation of human genes in the IFN system, we compare the representation of parametric features of human genes with different Log_2_(Fold Change) in experiments on human fibroblast cells stimulated with IFNs (Log_2_(Fold Change) > 0). Two features on the co-occurrence of SLims are not taken into consideration here as they are more subjective than the other parametric features and are greatly influenced by the number of focused SLims. Upon the calculation of PCC and the result of hypothesis tests, we find 168 features highly associated with the level of IFN-triggered stimulations (Student t-tests: *p* < 0.05) (**S3 Data**). Among them, 118 features show a positive correlation (Fig 10) while the remaining 50 features show a negative correlation (Fig 11) with the change of up-regulation in IFN experiments. Three features including the number of ORF, alternative splicing results, and exons in the canonical transcripts are encoded from characteristics of the gene. Two features, i.e., average dN/dS and average dS within human paralogues are encoded based on the sequence alignment results from the Ensembl [31]. 140 and 22 features are encoded from the genetic sequence and proteomic sequence respectively. The last one, i.e., neighbourhood connectivity is obtained from the network profile of a human interactome constructed based on experimentally verified data in the HIPPIE [47].

**Fig 10.**
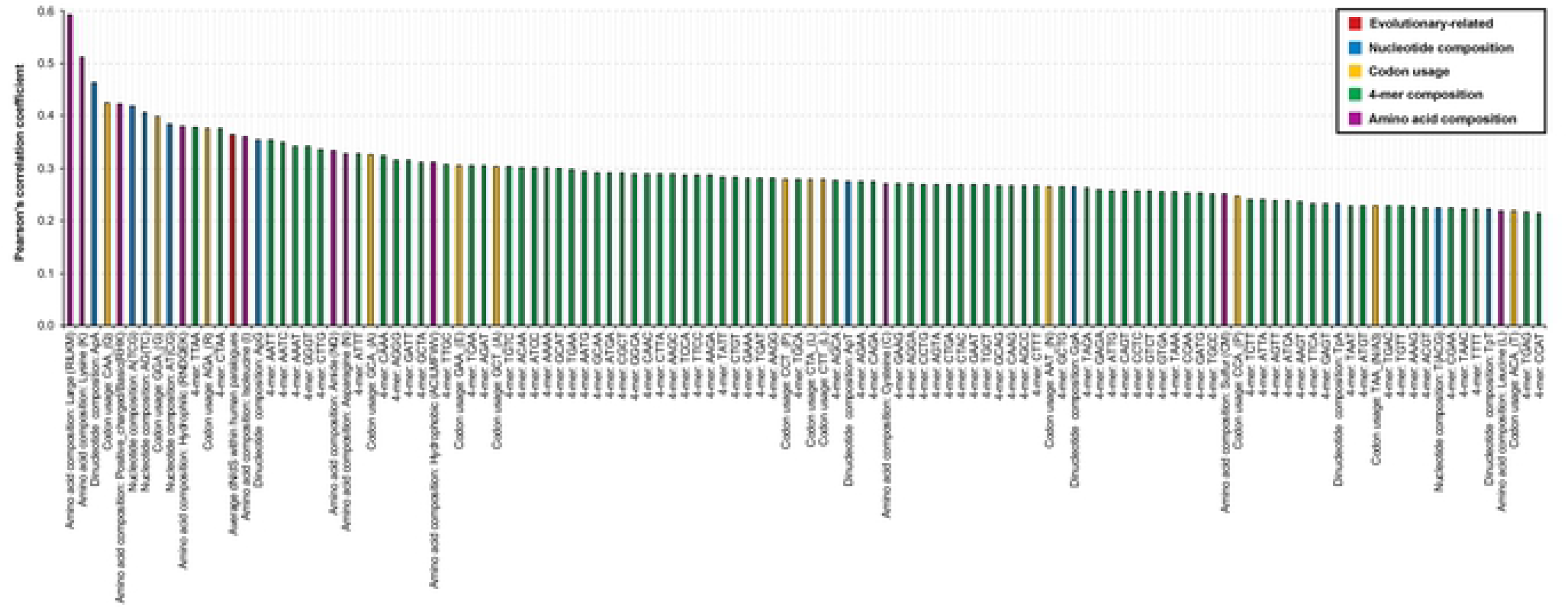
118 features positively associated with higher up-regulation after IFN treatments in human fibroblast cells (Student t-tests: p < 0.05). Detailed results about PCC and hypothesis tests are provided in **S3 Data**. Abbreviations: IFNs, interferons; PCC, Pearson’s correlation coefficient; dN, non-synonymous substitutions per non-synonymous site; dS synonymous substitutions per synonymous site.

**Fig 11.**
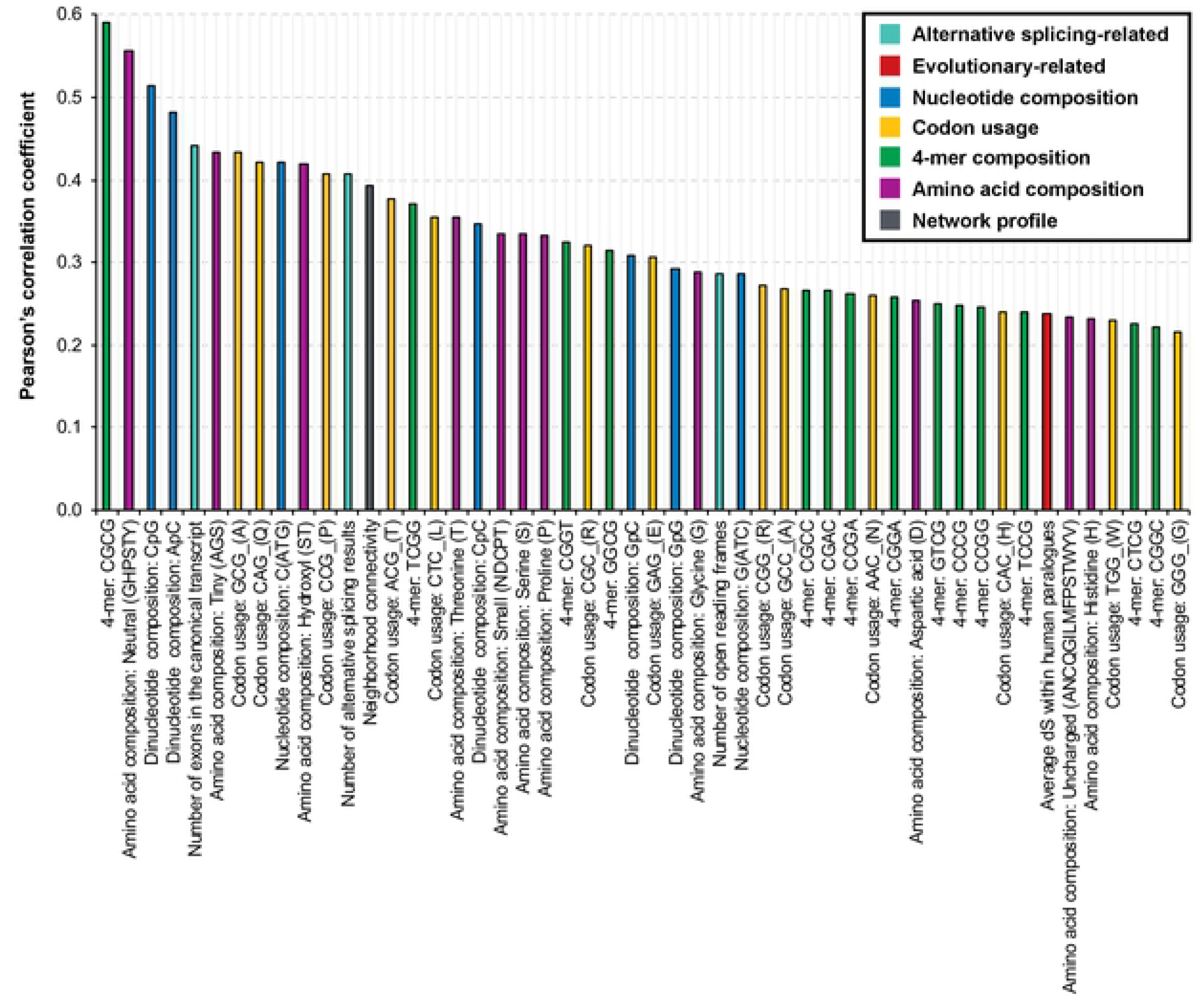
50 features negatively associated with higher up-regulation after IFN treatments in human fibroblast cells (Student t-tests: p < 0.05). Detailed results about PCC and hypothesis tests are provided in **S3 Data**. Abbreviations: IFNs, interferons; PCC, Pearson’s correlation coefficient; dS, synonymous substitutions per synonymous site.

In the positive group, the feature of ‘large’ amino acid compositions that includes the composition of five amino acids with geometric volume ranged from 163 to 173 cubic angstroms is ranked the first for having the highest PCC at 0.593 (Student t-test: *p* = 2.8E-09). This feature was not highlighted previously as it did not have a strong signal for discriminating ISGs from non-ISGs (Mann-Whitney U test: *p* > 0.05). Similar phenomena can be found on 87 features (64 positive correlations and 23 negative correlations) such as AG-content, ApG content and previously mentioned neutral amino acid composition. The strongest negative correlation between feature representation and IFN-triggered stimulations is found on the feature of 4-mer ‘CGCG’ (PCC = −0.593, *p* = 3.2E-09). This feature also shows a differential distribution between ISGs and non-ISGs, which provides useful information to distinguish ISGs from non-ISGs. Similar phenomena can be found on 81 features (54 positive correlations and 27 negative correlations) such as previously mentioned GC-content, CpG content and the usage of codon ‘GCG’ coding for amino acid ‘A’. Collectively, the biased effect on the basic composition of nucleotides influences the correlation between the representation of sequence-based features and IFN-triggered stimulations. Human genes that show over-representation in more features listed in Fig 10 are expected to be more up-regulated after type-I IFN treatments at least in human fibroblast cells. Meanwhile, the under-representation of features listed in Fig 11 will also contribute to the level of up-regulation in the IFN experiments.

### Difference in feature representation of interferon-repressed genes and genes with low levels of expression

We group human genes into two classes based on their response to the type I IFNs in human fibroblast cells. Genes significantly up-regulated in the IFN experiments are included in the ISG class, while those that do not are put into the non-ISG class. However, there is also another group of genes down-regulated in the presence of IFNs, i.e., IRGs. They are labelled as non-ISGs, but contain unique patterns that constitute an important aspect of the IFN response [8]. Some of these IRGs are not up-regulated in any known type I IFN systems, thus have been placed in a refined non-ISGs class for analyses and predictions. Additionally, there are a number of genes that have insufficient levels of expression in the experiments to determine a fold change. Here, we use the previously defined features to compare ISG with IRGs and ELGs.

As shown in Fig 12, IRGs are differentially represented to a lower extent in the majority of nucleotide 4-mer compositions than ISGs, which indicates the deficiency of some nucleotide sequence patterns in the coding region of IRGs. Note that, many nucleotide 4-mer composition features are more suppressed in ISGs than non-ISGs although the differences are small. The biased representation of these features in IRGs suggests that IRGs have characteristics similar to ISGs rather than non-ISGs. Additionally, there are a very limited number of features relating to evolutionary conservation, nucleotide compositions or codon usages showing obvious differences between ISGs and IRGs, but many of them are differentially represented when comparing ISGs with non-ISGs. Therefore, involving IRGs in the class of non-ISGs will increase the risk for machine learning models to produce more false positives. However, there are some informative features differentiating IRGs from ISGs. For example, comparing with ISGs, IRGs are more enriched in CpGs (Mann-Whitney U test: *p* = 5.6E-03), which is also mentioned in [70]. IRGs tend to have higher closeness centrality and neighbourhood connectivity than ISGs (Mann-Whitney U test: *p* = 0.04 and 6.4E-06 respectively), suggesting IRGs tend to be closer to the centre of the human PPI network and connected to key proteins with many interaction partners. Differences in some amino acid composition features between ISGs and IRGs can also be observed. Therefore, good predictability is still expected when using features extracted from proteins sequences.

**Fig 12.**
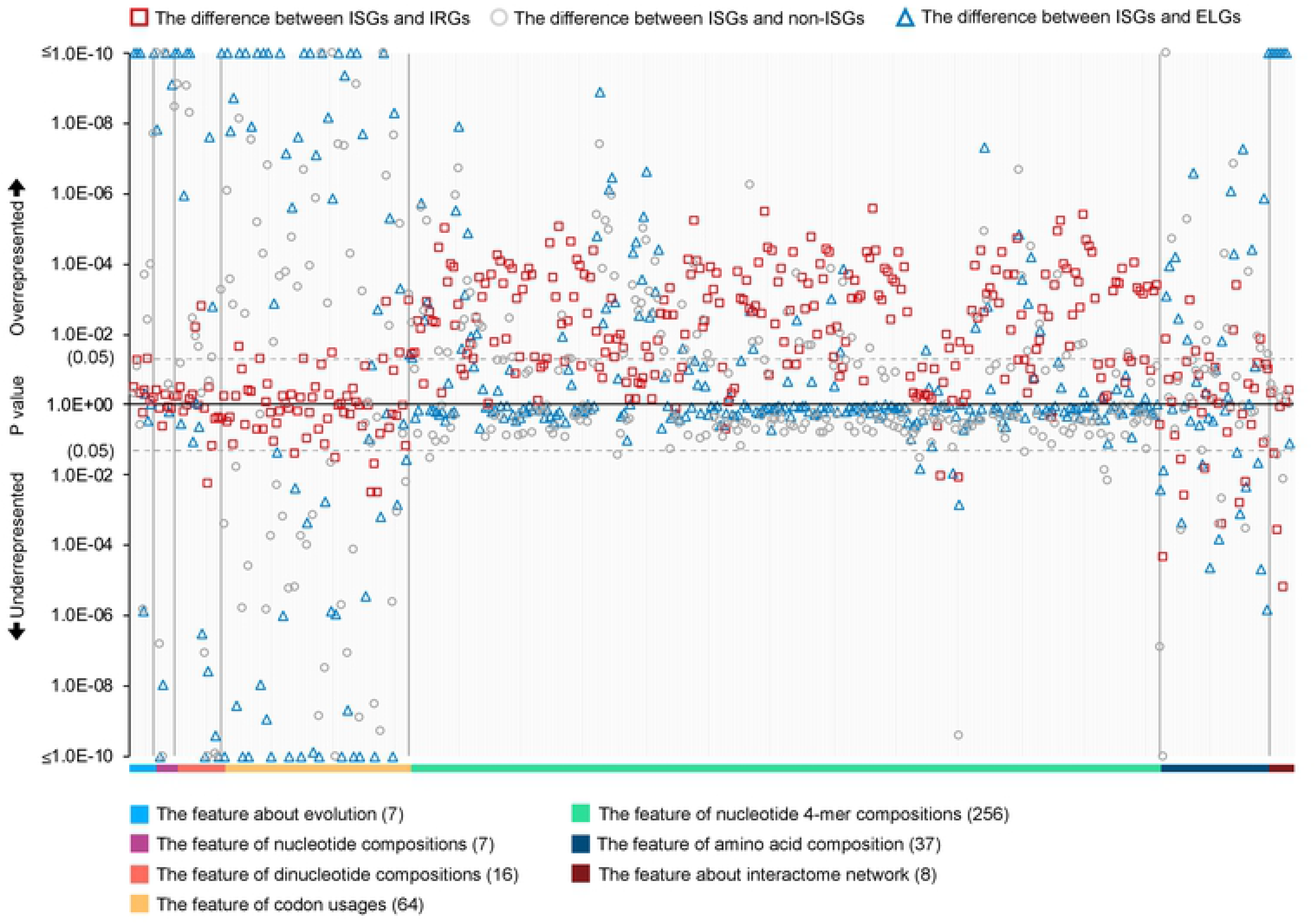
Differential expressions of parametric features between different genes and their coded proteins. Mann-Whitney U tests are applied for hypothesis testing and the results were provided in the **S2 Data**. Here, ISGs and non-ISGs are taken from dataset S2; IRGs and ELGs are taken from dataset S1; the background human genes are from dataset S1. Abbreviations: ISGs, interferon-upregulated genes; IRGs, interferon-repressed genes; non-ISGs, human genes not significantly up-regulated by interferons; ELGs, expression-limited human genes in IFN experiments.

Fig 12 illustrates 161 features showing significant differences (Mann-Whitney U tests: *p* < 0.05) in the representation of ISGs and ELGs. An estimated 82% of these features are also differentially represented between ISGs and non-ISGs. 79% of these significant features show similar over-representation or under-representation in two comparisons, i.e., ISGs versus ELGs and ISGs versus non-ISGs. These ratios indicate that the majority of ELGs are less likely to be ISGs based on their feature profile as well as their low expression levels in cells induced with IFNs. Network analyses show that ELG products tend to have lower values of all calculated network features with the exception of topological coefficient than ISG products, suggesting that they are less connected by other human proteins in the human PPI network. Particularly, their abnormal representation on the feature of average shortest paths indicating that some ELGs may still have high connectivity in the human PPI network, e.g., vascular cell adhesion molecule 1 (VCAM1) and ubiquitin D (UBD).

### Implementation with machine learning framework

In this study, we encode 397 parametric and 7046 non-parametric features for the analyses. As an excess of features will greatly increase the dimension of feature spaces and complicate the classification task for SVM [53], we limit the number of SLim_DNAs to the top 100 based on the adjusted p-value and we expect these to be sufficient to provide a picture of SLim patterns in the coding region of the canonical transcript. Accordingly, features measuring the co-occurrence status of multiple SLim_DNAs are recalculated based on the selected 100 SLim_DNAs. To reduce the impact of noisy data toward classifications, we only use the refined ISGs and non-ISGs, i.e., dataset S2 in machine learning.

Measured by SN, SP, MCC and AUC, the initial prediction results shown in Table 4 indicate that proteome-based features, including those deciphered from protein sequences and the human interactome, perform much better than genome-based features presumably due to overfitting of the model [71]. Using parametric features that take advantage of both genetic and proteomic aspects shows a good improvement in tests. The non-parametric features used in this study give a binary statement for the occurrence of SLims in genetic and proteomic sequences but seem not to perform well and disrupt the model when they are combined with parametric features. The results shown in the previous analyses also indicate that there are a considerable number of disruptive features hidden in the set (Fig. 5, Fig 7**, and** Fig 9). The similar attributes of ISGs and IRGs (Fig. 12) lead to lots of noisy data biasing the classifiers. This situation is not ameliorated and becomes more difficult when using other machine learning algorithms such as k-nearest neighbors (KNN), decision tree (DT), random forest (RF) (Table 4) [72, 73]. As some genes respond to IFNs in a cell-specific manner [2], it is hard to produce predictions unless we detect key discriminating features, which are robust to the change of biological environment.

**Table 4.**
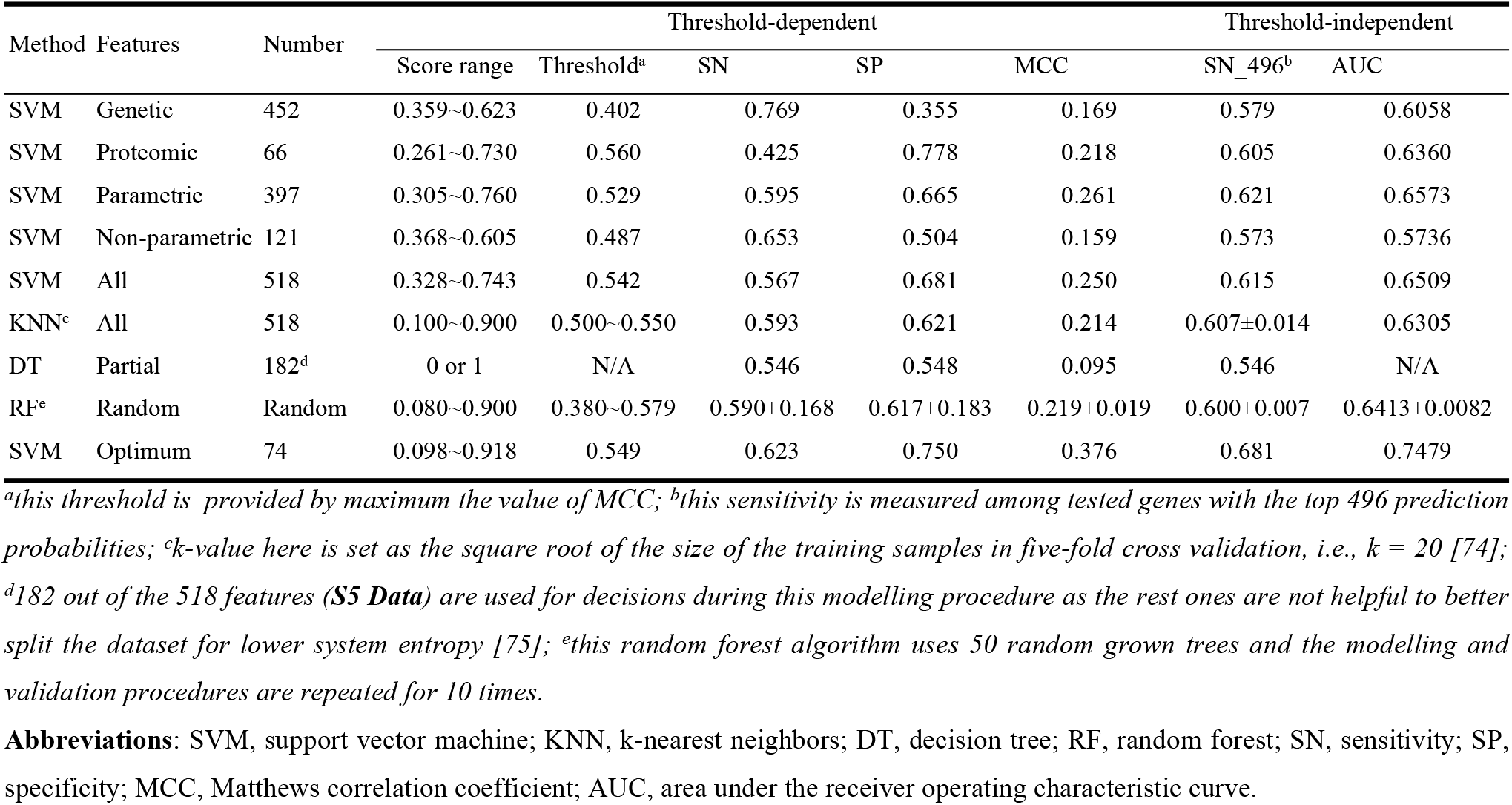
The performance of different feature combinations on the training dataset S2’ via five-fold cross validation.

| Method | Features | Number | Threshold-dependent |  |  |  | Threshold-independent |  |  |
| --- | --- | --- | --- | --- | --- | --- | --- | --- | --- |
|  |  |  | Score range | Threshold<sup>a</sup> | SN | SP | MCC | SN<sub>496</sub><sup>b</sup> | AUC |
| SVM | Genetic | 452 | 0.359~0.623 | 0.402 | 0.769 | 0.355 | 0.169 | 0.579 | 0.6058 |
| SVM | Proteomic | 66 | 0.261~0.730 | 0.560 | 0.425 | 0.778 | 0.218 | 0.605 | 0.6360 |
| SVM | Parametric | 397 | 0.305~0.760 | 0.529 | 0.595 | 0.665 | 0.261 | 0.621 | 0.6573 |
| SVM | Non-parametric | 121 | 0.368~0.605 | 0.487 | 0.653 | 0.504 | 0.159 | 0.573 | 0.5736 |
| SVM | All | 518 | 0.328~0.743 | 0.542 | 0.567 | 0.681 | 0.250 | 0.615 | 0.6509 |
| KNN<sup>c</sup> | All | 518 | 0.100~0.900 | 0.500~0.550 | 0.593 | 0.621 | 0.214 | 0.607±0.014 | 0.6305 |
| DT | Partial | 182<sup>d</sup> | 0 or 1 | N/A | 0.546 | 0.548 | 0.095 | 0.546 | N/A |
| RF<sup>e</sup> | Random | Random | 0.080~0.900 | 0.380~0.579 | 0.590±0.168 | 0.617±0.183 | 0.219±0.019 | 0.600±0.007 | 0.6413±0.0082 |
| SVM | Optimum | 74 | 0.098~0.918 | 0.549 | 0.623 | 0.750 | 0.376 | 0.681 | 0.7479 |

<sup>a</sup>this threshold is provided by maximum the value of MCC; <sup>b</sup>this sensitivity is measured among tested genes with the top 496 prediction probabilities; <sup>c</sup>k-value here is set as the square root of the size of the training samples in five-fold cross validation, i.e., $k = 20$ [74];
<sup>d</sup>182 out of the 518 features (**S5 Data**) are used for decisions during this modelling procedure as the rest ones are not helpful to better split the dataset for lower system entropy [75]; <sup>e</sup>this random forest algorithm uses 50 random grown trees and the modelling and validation procedures are repeated for 10 times.
**Abbreviations:** SVM, support vector machine; KNN, k-nearest neighbors; DT, decision tree; RF, random forest; SN, sensitivity; SP, specificity; MCC, Matthews correlation coefficient; AUC, area under the receiver operating characteristic curve.

Considering these drawbacks, we design an AUC-driven subtractive iteration algorithm (ASI) (Fig 2) to remove as many disruptive features as possible (Fig 13A). Pre-processing using the ASI algorithm shows that there are at least 28% of bad features disrupting the prediction model. They include 34% of features on codon usages and 50% of SLim features, thus, explaining the poor performance of the model trained with non-parametric features (Table 4). However, the loss of some of the individual nucleotide 4-mer feature seems not to influence the performance of the classifier at this stage, but the similarities between IRGs and ISGs (Fig 12) particularly in these 4-mer features is a cause for concern when the model is used to predict new data especially unknown IRGs. When using the ASI algorithm, the number of disrupting features does not stabilise and until the algorithm reaches the 11-th iterations when the number of disrupting features becomes zero. The remaining 74 features constitute our optimum feature set for the prediction of ISGs (Table 5). Among them, 14 and 9 features have positive and negative correlations with the level of up-regulation in IFN experiments. During the procedure, the AUC keeps increasing and reaches 0.7479 after 11 iterations. The MCC also shows an overall improvement although it fluctuates slightly during the last few iterations. By degressively ranking the probability calculated by the prediction model, we found 68.1% of the 496 genes (equal to the number of ISGs in the training dataset) are successfully predicted as ISGs. Fig 13B illustrates the distribution of probability scores generated by the ASI-optimised model for human genes with different expressions in IFN experiments. Human genes with higher up-regulation in IFN experiments tend to obtain higher probability score from our optimised machine learning model (PCC = 0.243, *p* = 4.2E-10). However, there are also some ISGs incorrectly predicted by our model even though they are highly up-regulated, e.g., basic leucine zipper ATF-like transcription factor 2 (BATF2, probability score = 0.34). The model produces 33 ISGs with a probability score higher than 0.8 but such figure for non-ISGs reduces to six, including one IRG, i.e., tripartite motif containing 59 (TRIM59). The highest probability score within non-ISGs was found on ubiquitin conjugating enzyme E2 R2 (UBE2R2, probability score = 0.88). It contains many features similar to ISGs but is not differentially expressed in the presence of IFN in fibroblast cells [8]. The lowest probability score within ISGs is found on cap methyltransferase 1 (CMTR1, probability score = 0.12) due to the weak signal from its features. For example, CMTR1 protein does not contain any ISG-favoured SLim_AA listed in Table 2. The influence of IRGs on the prediction is reflected in the training dataset but is not significant. Compared with human genes not differentially expressed in the IFN experiments, i.e., non-ISGs but not IRGs, there are slightly more IRGs unsuccessfully classified when using a threshold of 0.549 (Pearson’s chi-squared tests: *M_1_* = 27%, *M_2_* = 24%, *p* > 0.05).

**Fig 13.**
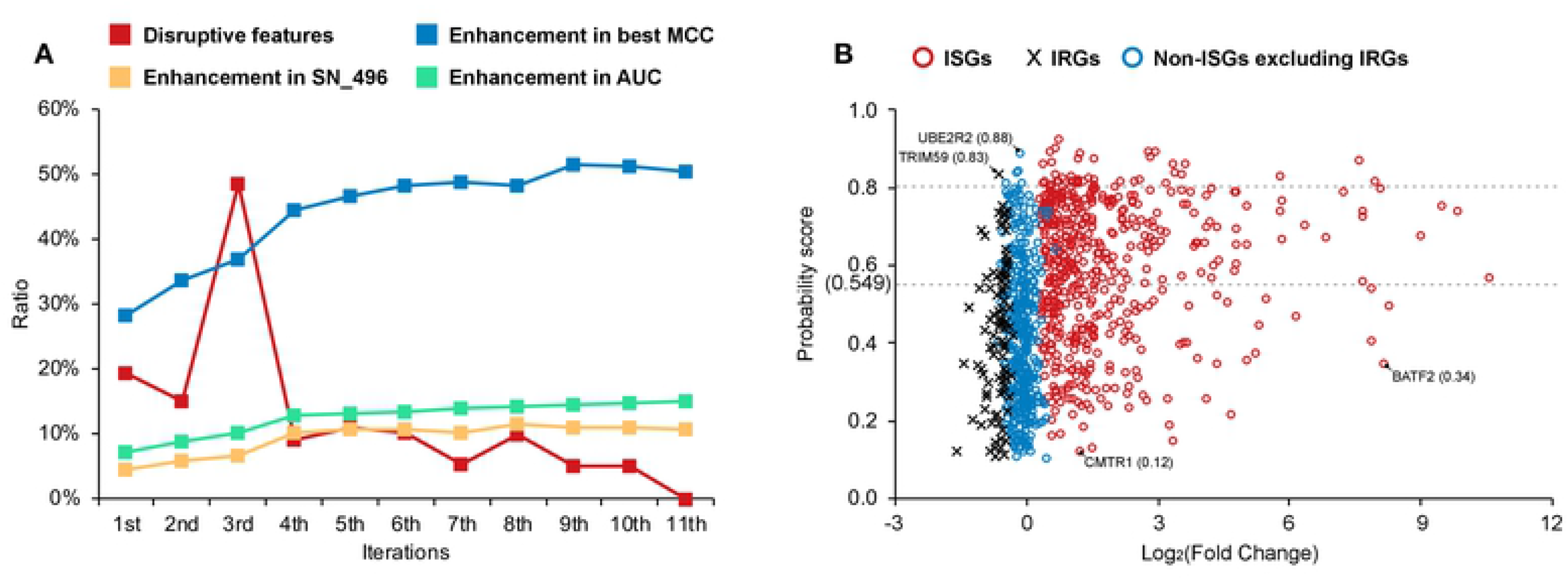
The optimisation on the machine learning model with the ASI algorithm. (A) shows the change of the prediction models based on the one generated with all 518 features (disruptive feature vector = 144, best MCC = 0.250, SN_496 = 0.615, and AUC = 0.6509). (B) shows the distribution of probability scores generated by the ASI-optimised model for human genes with different expression levels in the IFN system. ISGs and non-ISGs shown in (B) are randomly selected with an undersampling strategy on dataset S2. The list of gene names can be found in **S1 Data**. Abbreviations: SN, sensitivity; SN_496, sensitivity of predicted genes with the top 496 probability scores, MCC, Matthews correlation coefficient; AUC, area under the receiver operating characteristic curve; ASI, AUC-driven subtractive iteration algorithm; IFN, interferon, ISGs, interferons-stimulated genes; IRGs, interferon-repressed genes; non-ISGs, interferons-non-up-regulated genes; UBE2R2, ubiquitin conjugating enzyme E2 R2; TRIM59, tripartite motif containing 59; CMTR1, cap methyltransferase 1; BATF2, basic leucine zipper ATF-like transcription factor 2.

**Table 5.**
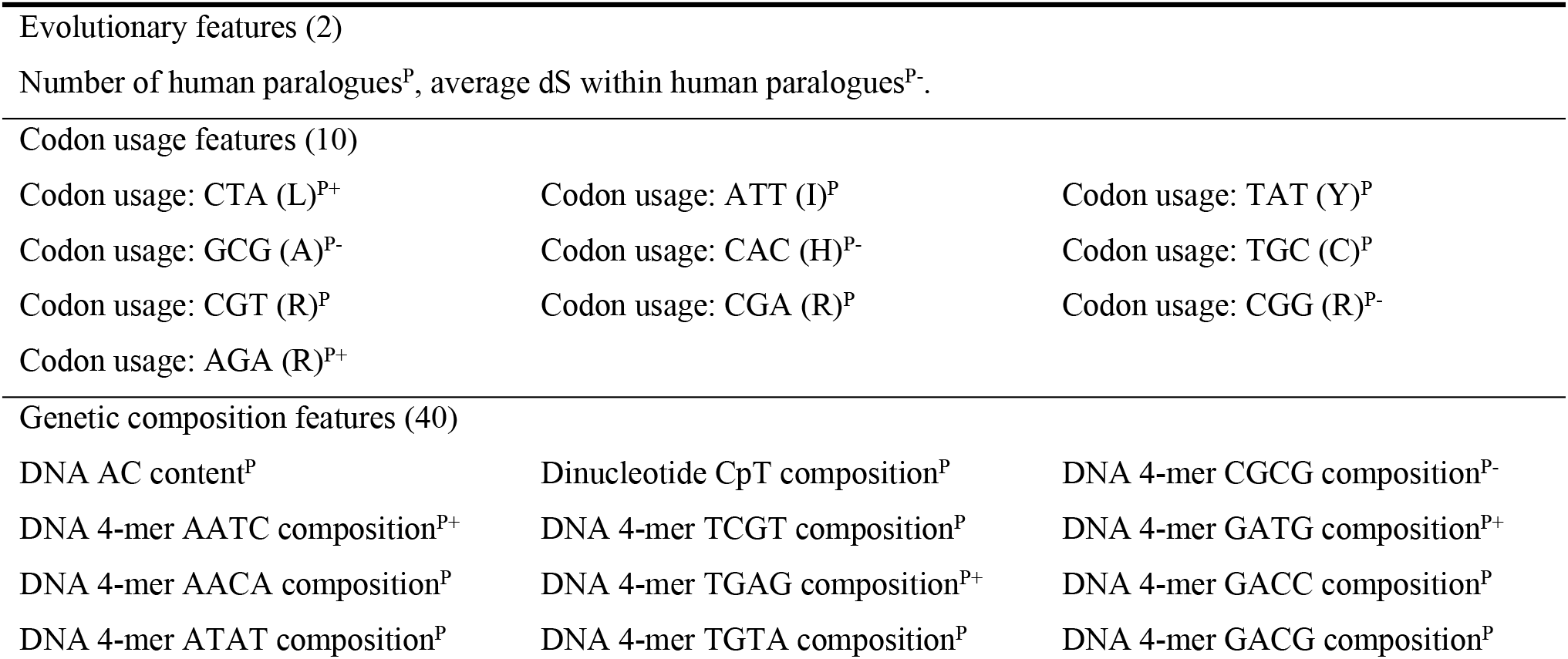
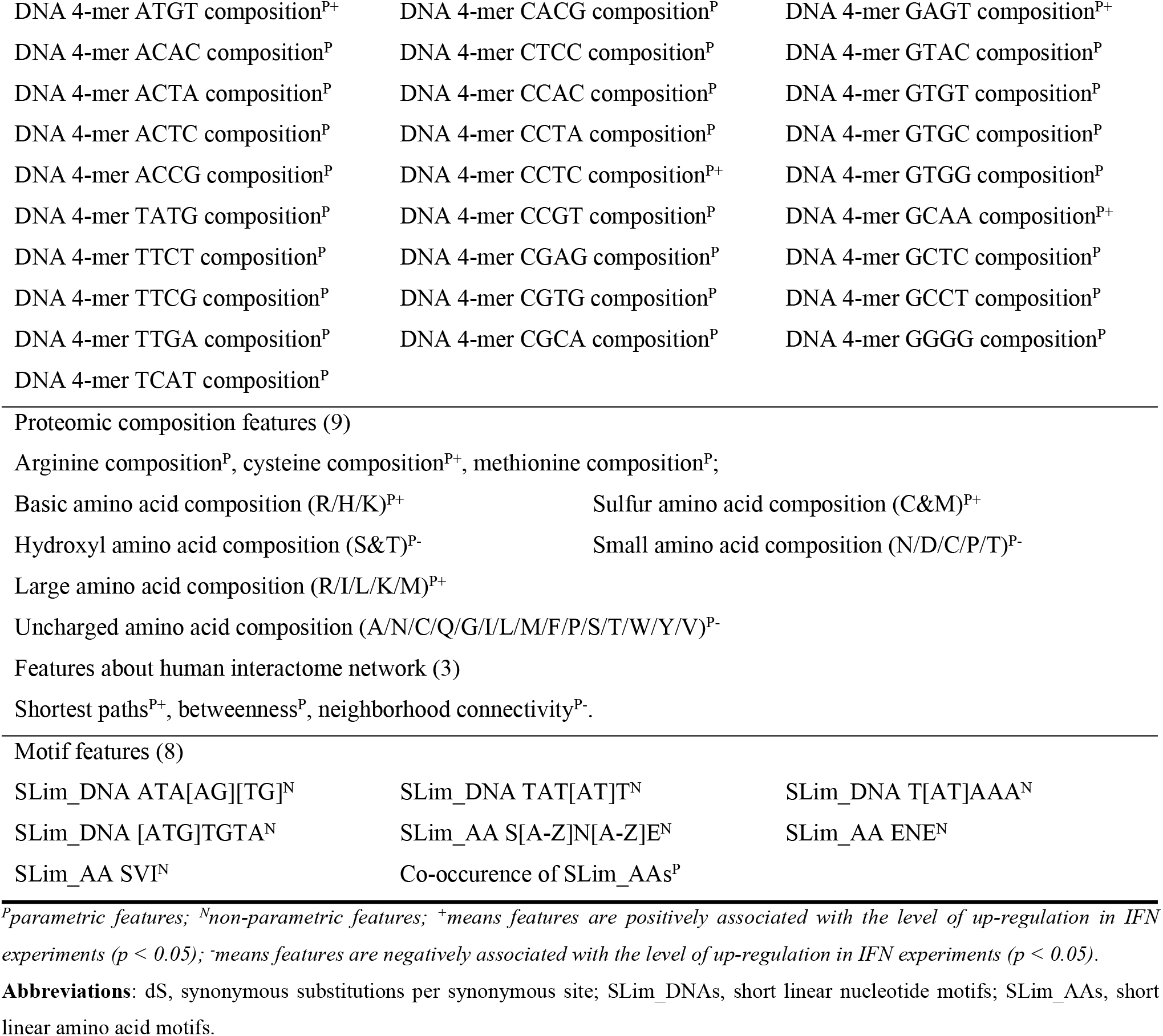
The optimum 74 features contributing to the prediction of ISGs.

|  |  |  |
| --- | --- | --- |
| Evolutionary features (2) |  |  |
| Number of human paralogues<sup>P</sup>, average dS within human paralogues<sup>P-</sup>. |  |  |
| Codon usage features (10) |  |  |
| Codon usage: CTA (L)<sup>P+</sup> | Codon usage: ATT (I)<sup>P</sup> | Codon usage: TAT (Y)<sup>P</sup> |
| Codon usage: GCG (A)<sup>P-</sup> | Codon usage: CAC (H)<sup>P-</sup> | Codon usage: TGC (C)<sup>P</sup> |
| Codon usage: CGT (R)<sup>P</sup> | Codon usage: CGA (R)<sup>P</sup> | Codon usage: CGG (R)<sup>P-</sup> |
| Codon usage: AGA (R)<sup>P+</sup> |  |  |
| Genetic composition features (40) |  |  |
| DNA AC content<sup>P</sup> | Dinucleotide CpT composition<sup>P</sup> | DNA 4-mer CGCG composition<sup>P-</sup> |
| DNA 4-mer AATC composition<sup>P+</sup> | DNA 4-mer TCGT composition<sup>P</sup> | DNA 4-mer GATG composition<sup>P+</sup> |
| DNA 4-mer AACA composition<sup>P</sup> | DNA 4-mer TGAG composition<sup>P+</sup> | DNA 4-mer GACC composition<sup>P</sup> |
| DNA 4-mer ATAT composition<sup>P</sup> | DNA 4-mer TGTA composition<sup>P</sup> | DNA 4-mer GACG composition<sup>P</sup> |
| DNA 4-mer ATGT composition<sup>P+</sup> | DNA 4-mer CACG composition<sup>P</sup> | DNA 4-mer GAGT composition<sup>P+</sup> |
| DNA 4-mer ACAC composition<sup>P</sup> | DNA 4-mer CTCC composition<sup>P</sup> | DNA 4-mer GTAC composition<sup>P</sup> |
| DNA 4-mer ACTA composition<sup>P</sup> | DNA 4-mer CCAC composition<sup>P</sup> | DNA 4-mer GTGT composition<sup>P</sup> |
| DNA 4-mer ACTC composition<sup>P</sup> | DNA 4-mer CCTA composition<sup>P</sup> | DNA 4-mer GTGC composition<sup>P</sup> |
| DNA 4-mer ACCG composition<sup>P</sup> | DNA 4-mer CCTC composition<sup>P+</sup> | DNA 4-mer GTGG composition<sup>P</sup> |
| DNA 4-mer TATG composition<sup>P</sup> | DNA 4-mer CCGT composition<sup>P</sup> | DNA 4-mer GCAA composition<sup>P+</sup> |
| DNA 4-mer TTCT composition<sup>P</sup> | DNA 4-mer CGAG composition<sup>P</sup> | DNA 4-mer GCTC composition<sup>P</sup> |
| DNA 4-mer TTCG composition<sup>P</sup> | DNA 4-mer CGTG composition<sup>P</sup> | DNA 4-mer GCCT composition<sup>P</sup> |
| DNA 4-mer TTGA composition<sup>P</sup> | DNA 4-mer CGCA composition<sup>P</sup> | DNA 4-mer GGGG composition<sup>P</sup> |
| DNA 4-mer TCAT composition<sup>P</sup> |  |  |
| Motif features (8) |  |  |
| SLim_DNA ATA[AG][TG]<sup>N</sup> | SLim_DNA TAT[AT]T<sup>N</sup> | SLim_DNA T[AT]AAA<sup>N</sup> |
| SLim_DNA [ATG]TGTA<sup>N</sup> | SLim_AA S[A-Z]N[A-Z]E<sup>N</sup> | SLim_AA ENE<sup>N</sup> |
| SLim_AA SVI<sup>N</sup> | Co-occurrence of SLim_AAs<sup>P</sup> |  |

<sup>P</sup>parametric features; <sup>N</sup>non-parametric features; <sup>+</sup>means features are positively associated with the level of up-regulation in IFN experiments ( $p < 0.05$ ); <sup>-</sup>means features are negatively associated with the level of up-regulation in IFN experiments ( $p < 0.05$ ).
**Abbreviations:** dS, synonymous substitutions per synonymous site; SLim\_DNAs, short linear nucleotide motifs; SLim\_AAs, short linear amino acid motifs.

### Review of different testing datasets

In this study, we train and optimise a SVM model from our training dataset, i.e., S2’, and prepare seven testing datasets to assess the generalisation capability of our model under different conditions. The S2’’ testing dataset is a subset of dataset S2. The prediction performance on this testing dataset is close to that in the training stage with an AUC of 0.7455 (Fig. 14). The best MCC value is achieved when setting the judgement threshold to 0.438, which means that the prediction model is sensitive to signals related to ISGs. In this case, it produces predictions with high sensitivity but inevitably produces many false positives, especially within the IRG class.

**Fig 14.**
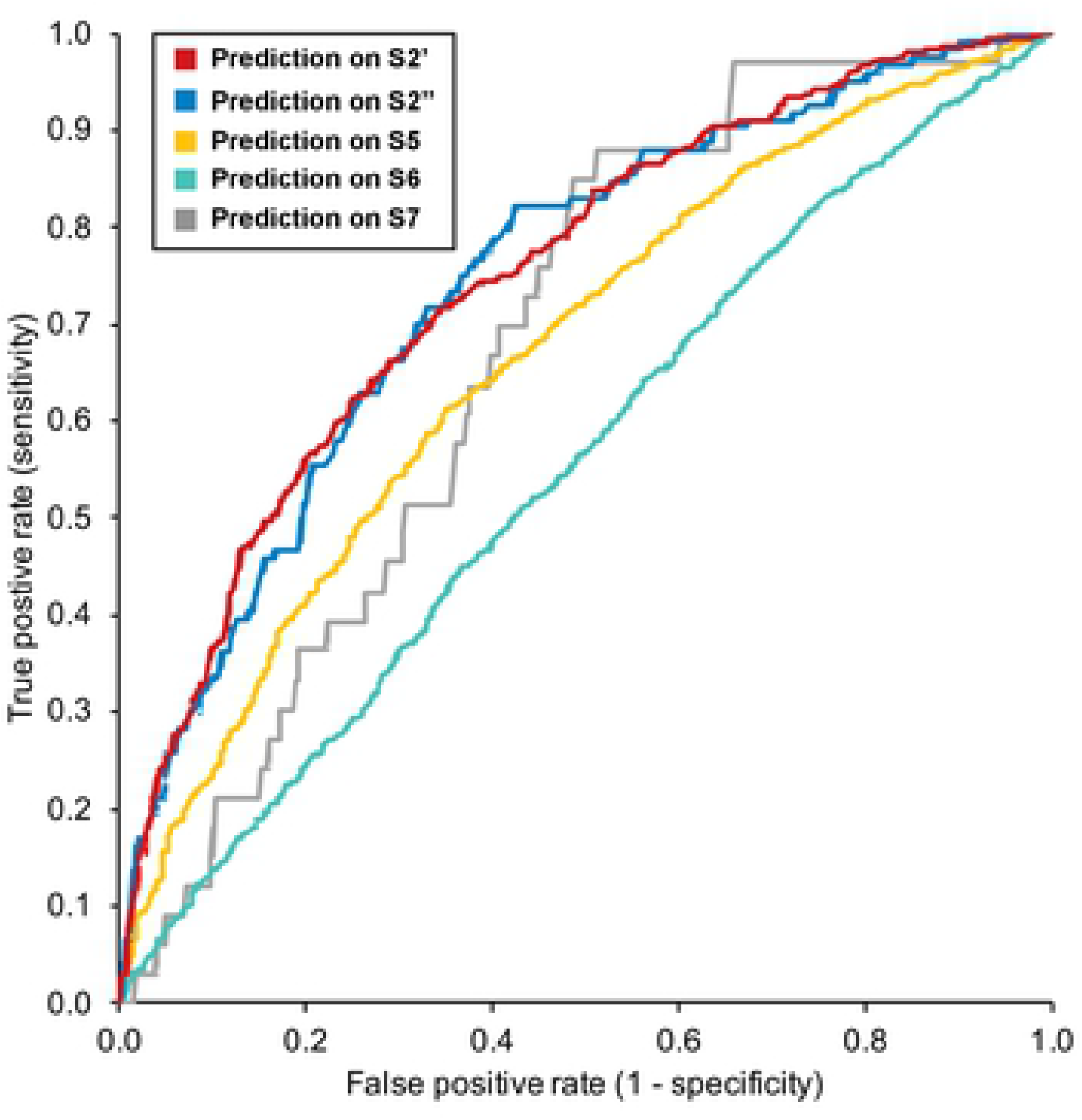
The performance of our optimised model on different datasets. S2’ is the training dataset used in this study. It randomly includes 496 ISGs and an equal number of non-ISGs from dataset S2 that contains ISGs/non-ISGs with high confidence (Table 1). Evaluation on this dataset in (A) is processed via five-fold cross validation. S2’’ is the testing dataset constructed with the remaining human genes in dataset S2. S5, S6, and S7 are collected from the Interferome database [21], including human genes with different responses to the type I, II and III IFNs, respectively. The label and usage of these human genes are provided in **S1 Data**. Abbreviations: AUC, area under the receiver operating characteristic curve; ISGs, interferon-stimulated genes; non-ISGs, human genes not significantly up-regulated by interferons.

In the S3 testing dataset, we use 695 ISGs with low confidence. The overall accuracy only reaches 44.0% when using a judgement threshold of 0.549, about 18% lower than SN under the same threshold in the training dataset S2’ (Table 4). This is expected as they have some inherent attributes that make them slightly up-regulated, silent or even repressed (e.g., become non-ISGs in other IFN systems) in response to some IFN-triggered signalling. On the other hand, on the S3 testing dataset, our machine learning model produces 38 (5.5%) ISGs with a probability score higher than 0.8. This number is also lower than on the training dataset S2’, which further indicates the relatively low confidence for ISGs included in testing dataset S3.

The S4 testing dataset is constructed to illustrate our hypothesis that there are some patterns shared among ISGs and IRGs at least in the type I IFN system in human fibroblast cells. On this testing dataset, the prediction accuracy is 60.2% under the judgement threshold of 0.549, about 15% lower than the SP under the same threshold in the training dataset S2’ (Table 4). Leucine rich repeat containing 2 (LRRC2), carbohydrate sulfotransferase 10 (CHST10) and eukaryotic translation elongation factor 1 epsilon 1 (EEF1E1) show strong signals of being ISGs (probability score > 0.9). In total, there are 56 (5.6%) IRGs being incorrectly predicted as ISGs with probability scores higher than 0.8. This high score is found in an estimated 8.1% of ISGs but is only observed in 1.2% of human genes not differentially expressed in the IFN experiments (Fig 13B). This result indicates that there is a considerable number of IRGs incorrectly predicted as ISGs in S4 testing dataset due to close distance to the ISGs in the high-dimensional feature space and this may be the case for any of the datasets. It also supports our hypothesis about the shared patterns from the machine learning aspect and is consistent with the results shown in Fig 12.

The next three testing datasets, i.e., S5, S6, and S7 are collected from the Interferome database [21] to test the applicability of the machine learning model across different IFN types. The ISGs in these testing datasets are all highly up-regulated (Log_2_(Fold Change) > 1.0) in the corresponding IFN systems while all the non-ISGs are not up-regulated after corresponding IFN treatments (Log_2_(Fold Change) < 0). The results shown in Fig 14 reveals that ISGs triggered by type I or III IFN signalling can still be predicted by our machine learning model, but the performance is limited (AUC = 0.6677 and 0.6754 respectively). However, it is almost impossible to make normal predictions with the current feature space for human genes up-regulated by type II IFNs (AUC = 0.5532).

The S8 testing dataset consists of 2217 human genes that are insufficiently expressed in the experiments in human fibroblast cells [8]. The results show that there are around 41.2% ELGs being predicted as ISGs when using a judgement threshold of 0.549. This is approximately 21% lower than the SN under the same threshold in the training dataset S2’ (Table 4). This suggests that there are more non-ISGs than ISGs in this dataset, which is consistent with the results of Fig 12. We find 10 ELGs with probability scores higher than 0.900: CD48 molecule, CD53 molecule, lipocalin 2 (LCN2), uncoupling protein 1 (UCP1), coiled-coil domain containing 68 (CCDC68), potassium calcium-activated channel subfamily M regulatory beta subunit 2 (KCNMB2), potassium voltage-gated channel interacting protein 4 (KCNIP4), zinc finger HIT-type containing 3 (ZNHIT3), serpin family B member 4 (SERPINB4), and fibrinogen silencer binding protein (FSBP). By retrieving data from the Genotype-Tissue Expression project [76], we find the expression of these ELGs are generally limited with the exception of CD53 and ZNHIT3 (Fig 15). The expression data of CD53 is not included in the OCISG database [8] and are also limited in the Interferome database [21]. It only shows slight up-regulation after type I treatments in blood, liver, and brain but there is currently no record of its expression level in the presence of type I IFNs in human fibroblast cells. ZNHIT3 is another well-expressed gene lacking information in the OCISG. In the Interferome databases, we find that ZNHIT3 can be up-regulated after IFN treatments in some fibroblast cells on skin. As for the remaining eight ELGs, despite their limited expression in human fibroblast cells, their features suggest that they are very likely to be IFN-stimulated in a currently untested cell type.

**Fig 15.**
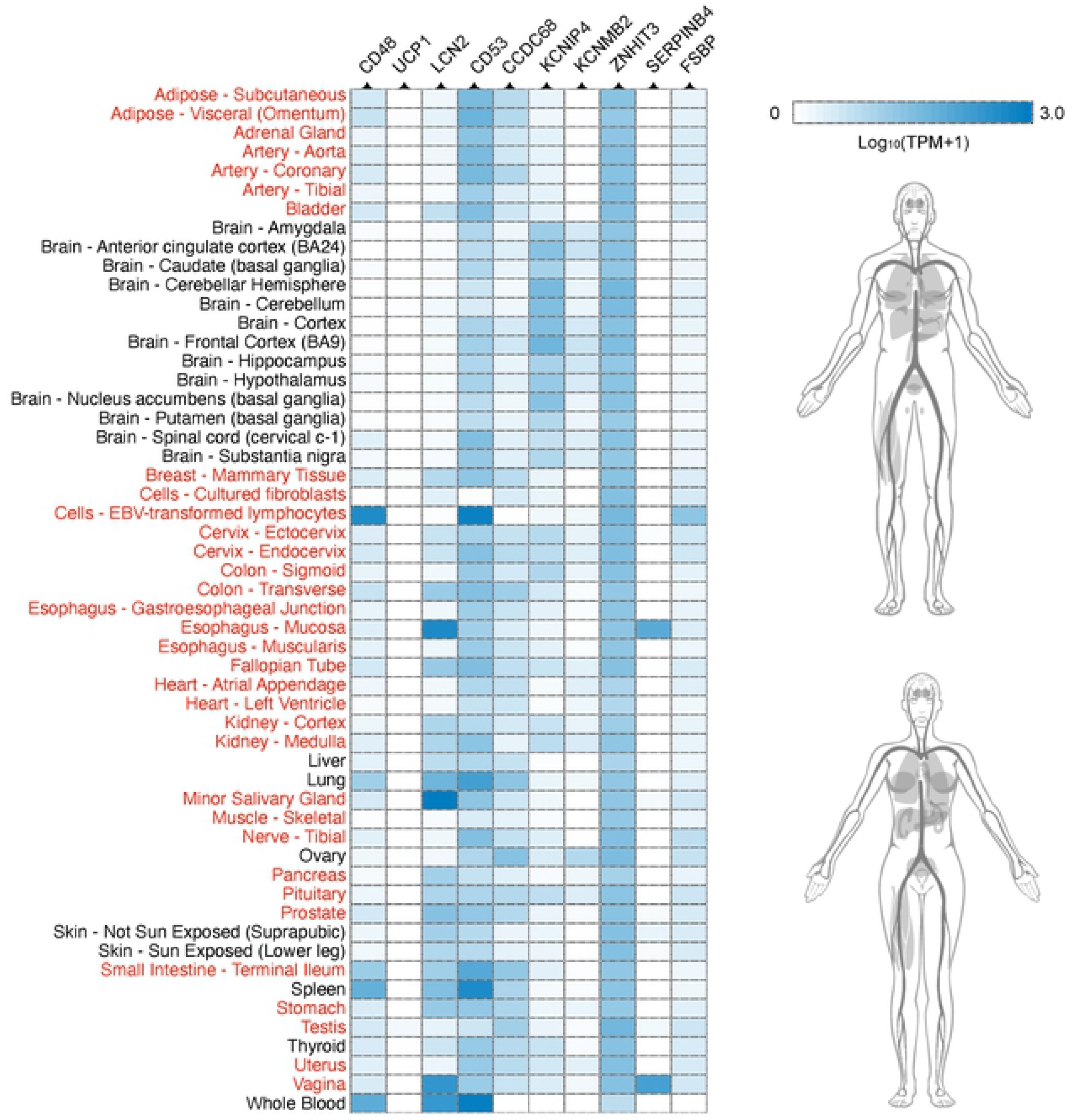
Expression of ELGs in different tissues. Expression data for ten ELGs are collected from the Genotype-Tissue Expression project (https://gtexportal.org/) [76]. The tissues in red are not included in the Interferome database [21]. White boxes in the heatmap indicate that there is no data available for genes in the corresponding tissues. The overall expression level of these ten ELGs are reflected via human perspective photo retrieved from Expression Atlas (https://www.ebi.ac.uk/gxa) [77]. Abbreviations: ELGs, human genes with limited expression in interferon experiments; TPM, transcripts per million; BA, Brodmann area; EBV, Epstein-Barr virus; UCP1, uncoupling protein 1; LCN2, lipocalin 2; CCDC68, coiled-coil domain containing 68; KCNIP4, potassium voltage-gated channel interacting protein 4; KCNMB2, potassium calcium-activated channel subfamily M regulatory beta subunit 2; ZNHIT3, zinc finger HIT-type containing 3; SERPINB4, serpin family B member 4; FSBP, fibrinogen silencer binding protein.

## Discussion

In this study, we investigate the characteristics that influence the expression of human genes in type I IFN experiments. We compare ISGs and non-ISGs through multiple procedures to guarantee strong signals for ISGs and to avoid cell-specific influences that result in the lack of ISGs expression in certain cell types [2]. Even some highly up-regulated ISGs can become down-regulated when the biological conditions change, exemplified by the performance of C-X-C motif chemokine ligand 10 (CXCL10) on liver biopsies after IFN-α treatment. This refinement is necessary as the representation of features between ISGs and the background human genes show that many non-ISGs especially IRGs have similar feature patterns to ISGs (Fig 4-7, Fig 12).

Generally, ISGs are less evolutionarily conserved with more human paralogues than non-ISGs. They have specific nucleotide patterns exemplified by the depletion of GC-content and have a unique codon usage preference in coding proteins. There are a number of SLim_DNAs widely observed in the cDNA of ISGs which are relatively rare in non-ISGs (**S4 Data**). Likewise, there are also many SLim_AAs highlighted in the sequences of ISG products that are absent or rare in non-ISGs (Table 2). In the human PPI network, ISG products tend to have higher betweenness than background human protein, indicating their more frequent interruption of the shortest path (geodesic distance) between different nodes. Abnormal expression or knockout of these proteins will increase the diameter of the network and may lead to some lethal consequences that are not tolerated in signalling pathways [78–80]. These ISG specific patterns may be the result of the evolution of the innate immune system in vertebrates and could be adaptations to the cellular environment induced by interferon following a pathogenic infection [81]. It is also possible that some of the particular SLim_DNAs and SLim_AAs may be important functionally as the cell changes from non-infected to infected. Experimental evidence will be necessary to investigate this.

Some inherent properties of ISGs facilitate or elevate their expression after IFN treatments but may also be used by viruses to escape from IFN-mediated antiviral response [19]. For instance, the representation of dN shows a more significant difference than that of dS within human paralogues Higher dN/dS ratio positively correlated with gene up-regulation following IFN treatments, but this means the gene is less conserved with more non-synonymous or nonsense mutations, which can often be associated to inherited diseases and cancer [82]. It will also facilitate the virus to interfere with IFN signalling through the JAK-STAT pathway and inactivate downstream cellular factors involved in IFN signal transductions [19]. Arginine is under-represented in ISG products compared to non-ISG products As arginine is essential for the normal proliferation and maturation of human T cells [83], such depletion in ISG products may leave a risk of inhibiting T-cell function and potentially increased susceptibility to infections [84]. On the other hand, the special pattern of ISGs also promotes the representation of some features even if they are not well represented in nature, exemplified by the higher cysteine composition in ISGs. We hypothesize that it may be helpful to activate T-cell to regulate protein synthesis, proliferation and secretion of immunoregulatory cytokines [85, 86]. For example, there are also some features, e.g., methionine composition, not differentially represented between ISGs and non-ISGs that play important roles in IFN-mediated immune responses. There is evidence for the methionine content playing a role in the biosynthesis of S-Adenosylmethionine (SAM), which can improve interferon signalling in cell culture [87, 88].

As previously mentioned, there are similar patterns between the feature representation of ISGs and IRGs, which leads to the unclear boundary for ISGs and non-ISGs in the feature space. We find significant differences on the representation of features on evolutionary conservation (Fig 4) between ISGs and non-ISGs, but these become non-significant when comparing ISGs with IRGs. Similar phenomena are observed on many features deciphered from the canonical transcript, e.g., dinucleotide composition and codon usage features. We suggest that IRGs can be viewed as additional ISGs as they also regulate the activity of human genes in response to IFNs, only negatively. On the other hand, despite so many similarities between ISGs and IRGs, the separate classification of these genes is still possible. 4-mer compositions can be considered as the key features as most of them are differentially represented between ISGs and IRGs (Fig 12). Using proteomic features can also help to differentiate ISGs from IRGs but is not as good as using 4-mer features.

In the machine learning framework, we develop the ASI algorithm to remove disruptive features but keep features not influencing the prediction performance when being removed individually during iterations. Features may have synergistic effects thus the elimination of each feature leaves a different impact on the remaining ones even if these are individually useless for the improvement of the classifier. In this case, keeping as many useful features as possible seems to be a good option but will greatly increase the dimension of the feature space and the risk of overfitting [71]. By contrast, our ASI algorithm avoids such a risk and keeps the synergistic effect of different features through iterations.

In the prediction task, we find some previously labelled non-ISGs with very high probability scores, suggesting that they have many inherent properties enabling them to be stimulated after IFN treatments. Some of them, for example UBE2R2 has been shown to be significantly up-regulated after IFN-α treatment [89]. The non-ISG label was assigned because the relevant expression data in the presence of IFNs are not included in the OCISG [8] and the Interferome databases [21]. We also find ten ELGs with very high probability scores (> 0.9). Literature searches on these genes indicate that they are likely to be involved in the innate immune response and that their responses may be limited to certain tissues or cell types for which there is limited expression data in the Interferome database [21]. For example, LCN2 has been shown to mediate an innate immune response to bacterial infections by sequestering iron [90] and is induced in the central nervous system of mice infected with West Nile virus encephalitis [91]. CD48 was shown to increase in levels as a result of human IFN-α/β and human IFN-γ and these upregulate the expression of CD48 proteins at the surface of various cultured human cell lines [92]. Interestingly, CD48 is also the target of immune evasion by viruses [93] and has been captured in the genome of cytomegalovirus and undergone duplication [94]. Evidence for other ELGs is harder to assess, particularly those for which expression is absent in a range of tissues (e.g., UCP1 in Fig 15). UCP1 is a mitochondrial carrier protein expressed in brown adipose tissue (BAT) responsible for non-shivering thermogenesis [95]. It is possible that UCP1 is stimulated directly or indirectly by IFN in BAT resulting in the defended elevation of body temperature in response to infection. In this in silico study, we provide predictions for genes that show no basal expression in human fibroblasts but their stimulation by IFN and their role in immune defense requires testing experimentally.

The model developed in this study based on experimental data from human fibroblast cells stimulated by IFN can be generalised to type III systems, presumably because activations of type I and III ISGs are both controlled by ISRE [9] and aim to regulate host immune response [4–6]. However, our model cannot be applied to the prediction of type II ISGs (AUC = 0.5532), not only because of their different control elements, but because of their different roles in human immune activities (Fig 1) [10].

In summary, our analyses highlight some key sequence-based features that are helpful to distinguish ISGs from non-ISGs or IRGs. Our machine learning model is able to produce a list of putative ISGs to support IFN-related research. As knowledge of ISG functions continue to be elucidated by experimentalists, the *in-silico* approach applied here could in future be extended to classify the different functions of ISGs.

## Supporting information

**S1 Data. Basic information about human genes used in this study.**

(TXT)

**S2 Data. The result of Mann-Whitney U tests for parametric features.**

(TXT)

**S3 Data. Association between feature representations and IFN stimulations.**

(TXT)

**S4 Data. The result of Pearson’s chi-squared tests for sequence motifs.**

(TXT)

**S5 Data. Decision trees generated during five-cross validation on the training dataset S2’.**

(TXT)

## Acknowledgments

The authors wish to thank Prof Andrew Davison, Drs Suzannah Rihn and Sam Wilson for helpful discussions and recommendations, and Scott Arkison for help setting up the website.

## Author contributions

**Conceptualization:** David L. Robertson, Joseph Hughes, Quan Gu, Haiting Chai.

**Data curation:** Haiting Chai.

**Formal analysis:** Haiting Chai.

**Funding acquisition:** Haiting Chai, David L. Robertson.

**Webserver:** Haiting Chai.

**Supervision:** David L. Robertson, Joseph Hughes, Quan Gu.

**Writing-original draft:** Haiting Chai.

**Writing-review & editing:** David L. Robertson, Joseph Hughes, Quan Gu, Haiting Chai.

## Data availability statement

The implemented web server and data were freely accessible at http://isgpre.cvr.gla.ac.uk/.

